# Glycogen removal enhances ethylene production in *Synechocystis* sp. PCC 6803 but exposes a 2-oxoglutarate–GS-GOGAT–phosphate constraint

**DOI:** 10.64898/2026.10.25.761650

**Authors:** Tomáš Zavřel, Jacob Sebesta, Hakyung Lee, Tomáš Zlámal, Terezie Danišová, Roman Sándor, Martin Černý, Miroslav Berka, Martin Lukeš, Rune Höper, Ralf Steuer, Jan Červený, Jianping Yu

## Abstract

Cyanobacteria are promising platforms for converting CO_2_ directly into value-added chemicals. However, redirecting fixed carbon toward a product requires understanding of how cells balance the energetic and redox supplies and demands - and how efficiently metabolic flux can be channelled into product formation rather than being dissipated or excreted. Glycogen, synthesized *via* ADP-glucose pyrophosphorylase (GlgC), is the major native carbon and energy sink, and its removal forces cells to redirect excess carbon and reducing power into alternative pathways. As a target product we used ethylene, a major industrial building block currently derived almost entirely from fossil feedstocks, which can be produced photoautotrophically by the ethylene-forming enzyme (EFE) at the expense of 2-oxoglutarate (2-OG), a central node linking carbon and nitrogen metabolism. Here, we compared wild-type *Synechocystis* sp. PCC 6803 with strains carrying either a complete (*ΔglgC*) or attenuated, light-responsive (*LGS*; psbA2-driven *glgC*) glycogen synthesis block, with an EFE-expressing strain (*EFE*), and with the combined *ΔglgC+EFE* strain. All strains were cultivated across a light gradient (35–300 µmol photons m⁻² s⁻¹) under nitrogen-replete conditions. Integrating measurements of growth and ethylene production rates, chlorophyll-fluorescence analysis, quantitative proteomics, FTIR profiling, and overflow-metabolite analysis revealed that partial suppression of glycogen synthesis (*LGS*) enhanced growth under low irradiance but became detrimental above a critical light threshold. EFE expression in the *ΔglgC* background increased ethylene production in a light-dependent manner but further compromised growth. Both strains with EFE increased photosynthetic electron transport consistent with the carbon-sink effect, yet this electron surplus was not channelled proportionally into product or biomass. Instead, continuous 2-OG drain by EFE constrained flux through the GS-GOGAT node and induced a secondary inorganic phosphate limitation in *ΔglgC+EFE* that activated photoprotection and limited productivity. These results point to the 2-OG/GS-GOGAT/Pi axis as a key design constraint determining whether the surplus photosynthetic flux gets converted into product, dissipated through photoprotection, or released as overflow metabolites.

## 1. Introduction

Cyanobacteria convert carbon dioxide, water, and sunlight directly into biomass and a range of value-added compounds, making them attractive platforms for sustainable, carbon-neutral chemical production (Toepel et al., 2023). Realising this potential requires redirecting photosynthetically fixed carbon and reducing equivalents away from native sinks, toward a target product (Santos-Merino et al., 2021). This is challenging because cyanobacterial photosynthesis often operates below its maximal capacity and can be constrained by the availability of downstream metabolic sinks rather than by light capture or electron transport itself (Grund et al., 2019). Implementing a strong heterologous sink can therefore relieve this limitation and stimulate photosynthetic electron flow, an effect known as the “carbon-sink effect” (Grund et al., 2022). How efficiently the resulting surplus is channeled into product, however, depends on the capacity of the cell to accommodate the associated energetic, redox, and metabolic imbalances.

In cyanobacteria, glycogen is the major native carbon and energy sink. This glucose polymer is synthesised *via* ADP-glucose pyrophosphorylase, encoded by *glgC*, and serves as a metabolic buffer between carbon supply and demand. Glycogen stores surplus fixed carbon under energy-replete conditions and is remobilised through the oxidative pentose phosphate pathway to provide reducing power and sustain cellular metabolism during darkness (Cano et al., 2018; Shinde et al., 2020). Deletion of *glgC* removes this buffer and forces the cell to redirect excess carbon and reducing equivalents through alternative pathways. Accordingly, *ΔglgC* strains exhibit pronounced metabolic overflow, including excretion of organic acids such as 2-oxoglutarate (2-OG) and pyruvate (Carrieri et al., 2012; Gründel et al., 2012; Cano et al., 2018; Alford et al., 2025). This redistribution has been exploited to enhance photosynthetic production of organic acids and other heterologous products, including lactate and ethanol (van der Woude et al., 2014; Namakoshi et al., 2016).

Among potential target molecules, ethylene is particularly attractive: it is one of the most widely used building blocks of the chemical industry and is currently derived almost entirely from fossil feedstocks. Photoautotrophic ethylene production can be achieved by heterologous expression of the ethylene-forming enzyme (EFE), which couples the oxidative decarboxylation of 2-OG to arginine hydroxylation, producing ethylene together with CO_2_, succinate, guanidine, and pyrroline-5-carboxylate (Fukuda et al., 1992; Eckert et al., 2014). EFE therefore directly intersects a central node of cyanobacterial metabolism: 2-OG links carbon metabolism to nitrogen assimilation through the glutamine synthetase–glutamate synthase (GS-GOGAT) cycle and functions as a key signal in the PII regulatory network and for the master nitrogen regulator NtcA (Forchhammer and Selim, 2020). Combining *glgC* deletion with EFE expression is therefore appealing in principle: the carbon, ATP, and reducing power made available by reducing glycogen synthesis could potentially be channelled into ethylene production. However, the metabolic consequences of this combination are not straightforward. An earlier *ΔglgC+EFE* strain produced less ethylene than the corresponding wild-type (WT) background under nitrogen limitation despite elevated intracellular 2-OG level (Veetil et al., 2017). This finding demonstrates that increasing precursor availability does not necessarily translate into increased product formation and raises the question of how the resulting carbon and reducing-power surplus is redistributed.

Complete *glgC* deletion is associated with a severe growth penalty and obligate dependence on continuous light, limiting the utility of this genotype as a production chassis. An alternative strategy is to only reduce glycogen synthesis, such as in a low-glycogen strain (*LGS*) in which *glgC* expression is placed under the light-responsive *psbA2* promoter, allowing glycogen synthesis to be partially and dynamically restored under high irradiance (Carrieri et al., 2012; Cantrell et al., 2023). Such attenuation strategies can improve photosynthetic productivity while retaining some capacity for carbon storage. However, how partial and light-responsive suppression of glycogen synthesis reshapes photosynthetic electron transport, proteome allocation, and overflow metabolism — and how these responses compare with those caused by complete glycogen removal or EFE expression — remains poorly understood.

Here we systematically compare WT *Synechocystis* sp. PCC 6803 (hereafter *Synechocystis*) with strains carrying complete (*ΔglgC*) or attenuated (*LGS*) blocks in glycogen synthesis, with an EFE-expressing strain (*EFE*), and with the combined *ΔglgC+EFE* strain. We characterised these strains across a gradient of light intensities under nitrogen-replete, photoautotrophic conditions. By integrating growth and ethylene-production measurements with chlorophyll fluorescence-based assessment of photosynthetic electron transport and photoprotection, quantitative proteomics, FTIR-based analysis of macromolecular composition, and analysis of secreted overflow metabolites, we investigated how these interventions reshape the allocation of fixed carbon and reducing equivalents. We find that downregulating glycogen synthesis does not simply redirect the metabolic fluxes into biomass or product formation, but instead induces distinct metabolic and photoprotective responses. In the EFE-expressing background, continuous 2-OG consumption perturbs the GS-GOGAT node and reveals a secondary limitation in inorganic phosphate (Pi), further constraining productivity. We also identify distinct routes of carbon disposal, ranging from organic-acid overflow to intracellular metabolic rerouting. These results identify the 2-OG/GS-GOGAT/Pi axis as a central determinant of whether surplus flux is channelled into product, dissipated through photoprotection, or excreted as overflow, and show that removing a carbon-storage pathway does not merely redirect flux into a product but reconfigures whole-cell energy management. Framing these comparisons as an engineering problem, we address three questions highly relevant to the design of cyanobacterial production strains: (i) whether removing or attenuating the glycogen sink increases the fraction of fixed carbon and reducing equivalents reaching a heterologous product; (ii) which cellular processes capture or dissipate the resulting surplus; and (iii) what these constraints imply for the design of *Synechocystis* chassis carrying 2-OG-consuming pathways.

## 2. Materials and Methods

### 2.1 Strains generation

All experiments were performed using the model cyanobacterium *Synechocystis* sp. PCC 6803. Glucose-tolerant wild-type (WT) was used as a control strain. All heterologous strains were generated using homologous recombination and natural uptake transformation. *ΔglgC* and *LGS* were generated previously with *LGS* consisting of the *glgC* deletion complemented with *PpsbA2* promoter driving *glgC* expression (Carrieri et al., 2012). The *EFE* strain was previously generated and consists of the *PpsbA\** promoter, *RBSv32*, and a codon optimized *efe* sequence from *Pseudomonas syringae* pv *phaseolicola* PK2 (Wang et al., 2018). The *ΔglgC+EFE* strain was generated by transforming the *PpsbA*-RBS32-EFE* construct into the *glgC* knockout background. Deletions were confirmed by colony PCR. Correct *efe* sequences were verified by amplicon sequencing. *ΔglgC* was maintained with 50 µg mL^-1^ gentamicin, *LGS* with 50 µg mL^-1^ gentamicin and 50 µg mL^-1^ kanamycin, and EFE- containing strains included 50 µg mL^-1^ spectinomycin.

### 2.2 Cultivation conditions and growth rate monitoring

All strains were cultivated in 250 mL Erlenmeyer flasks containing approx. 150 mL of culture. Cultures were maintained in a controllable Percival cultivation chamber at 30 °C and 5% CO_2_ (v/v) in the atmosphere. Flasks were shaken at 150 rpm and illuminated with warm-white fluorescent lamps at photon flux densities (PFD) of 35 - 300 µmol photons m^-2^ s^-1^. Specific growth rates were determined from optical density at 720 nm (OD_720_), measured using an Infinite M2000 Pro plate reader (Tecan Group Ltd., Switzerland) in 96-well plates. Cultures were inoculated at an OD_720_ of 0.02–0.05 and acclimated to each tested light intensity for 24- 48 h before measurements.

### 2.3 Photosynthesis activity measurement

Photosynthetic performance was assessed using an AquaPen fluorometer (Photon System Instruments, Czechia), with data processing performed using the analysis pipeline available at https://www.cyano.tools/ (Zavřel, 2026). Three protocols were applied sequentially: fast fluorescence induction kinetics (OJIP), non-photochemical quenching (NPQ3) and rapid light curves (LC3). The following instrument settings were used for all measurements: measuring light (ML): 630 nm, flash pulse: 50%, super pulse: 50% (approximately 2 000 µmol photons m^-2^ s^-1^), and actinic pulse 100 µmol photons m^-2^ s^-1^. All other instrument settings were kept at their default values. Before measurement, cultures were diluted with fresh BG-11 medium to obtain a minimal fluorescence (F_O_) of 20 000 - 60 000 instrument units after dark acclimation. Samples were dark-acclimated for 15 min before measurement. For NPQ3 and LC3 measurements, cultures were continuously stirred at 300 rpm using a benchtop magnetic stirrer and a 5 x 3 mm magnetic bar. The following parameters were calculated from the NPQ3 measurements:

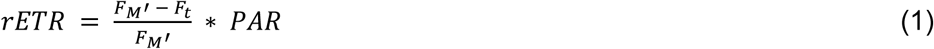

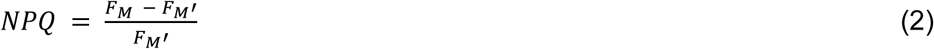

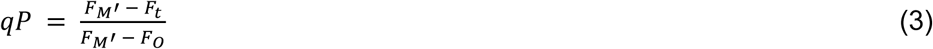

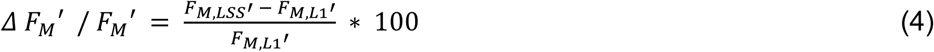

where rETR is the relative electron transport rate under actinic illumination, PAR is photosynthetically active radiation, NPQ is non-photochemical quenching, and qP is the coefficient of photochemical quenching (van Kooten and Snel, 1990). F₀ and F_M_ denote minimal and maximal fluorescence, respectively, in the dark-acclimated state, whereas F_t_ and

F_M_’ denote steady-state and maximal fluorescence, respectively, under actinic illumination. The subscripts L1 and Lss denote values measured at the first saturating pulse after the onset of actinic illumination and after full light acclimation, respectively, as designated by the AquaPen software (Fig. 2A-C). ΔF_M_’/F_M_’ describes the relative change in maximal fluorescence during light acclimation and was used as a proxy for the extent and direction of state transitions (Calzadilla and Kirilovsky, 2020; Kis et al., 2026), with positive values indicating a State 2→1 transition and negative values a State 1→2 transition. Parameters in Eqs. 1-3 were determined after full light acclimation (Lss).

**Fig. 1.**
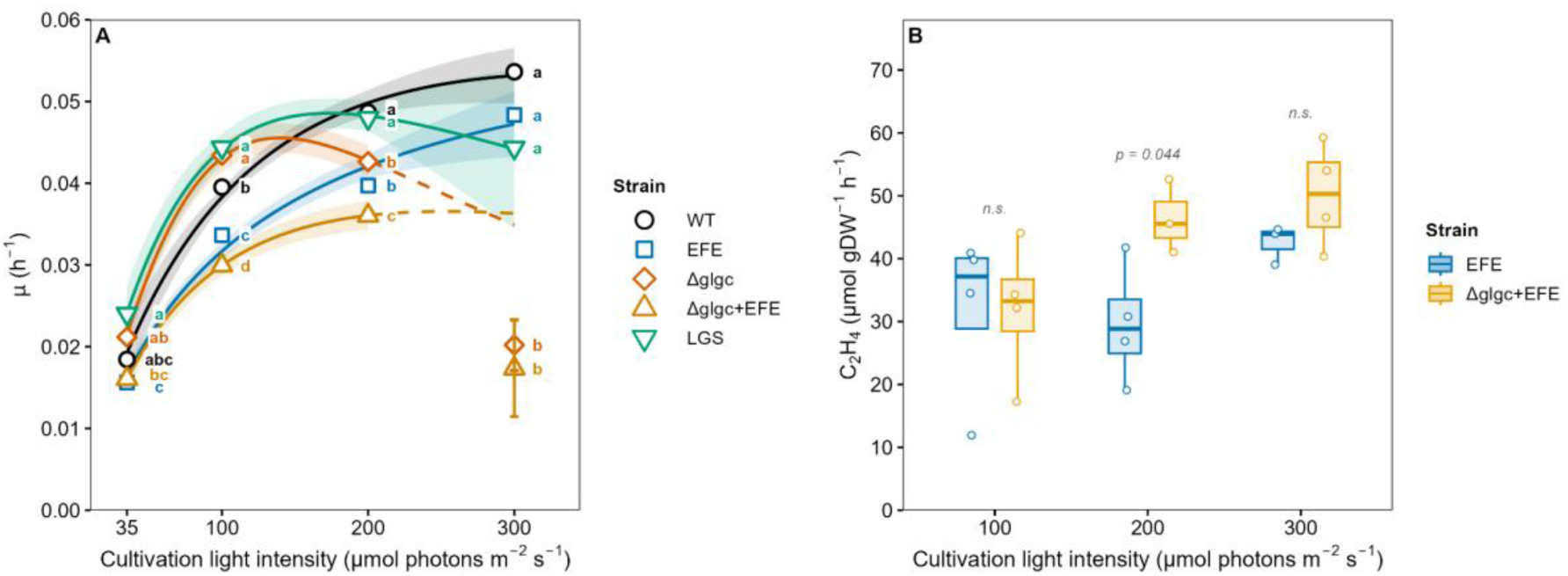
Specific growth rate of the tested strains (A) and ethylene production rates in strains expressing EFE (B). Growth-rate data in (A) were fitted with the Platt model (Platt et al., 1980) (solid lines) to obtain comparable light- response parameters; the model does not fully capture the threshold-like growth suppression observed in *ΔglgC* and *ΔglgC+EFE* at high irradiance (dashed lines). Letters indicate significant differences between strains within a single light intensity. The p-value is given where p < 0.05 and n.s. (not significant) otherwise; n = 3-4.

**Fig. 2.**
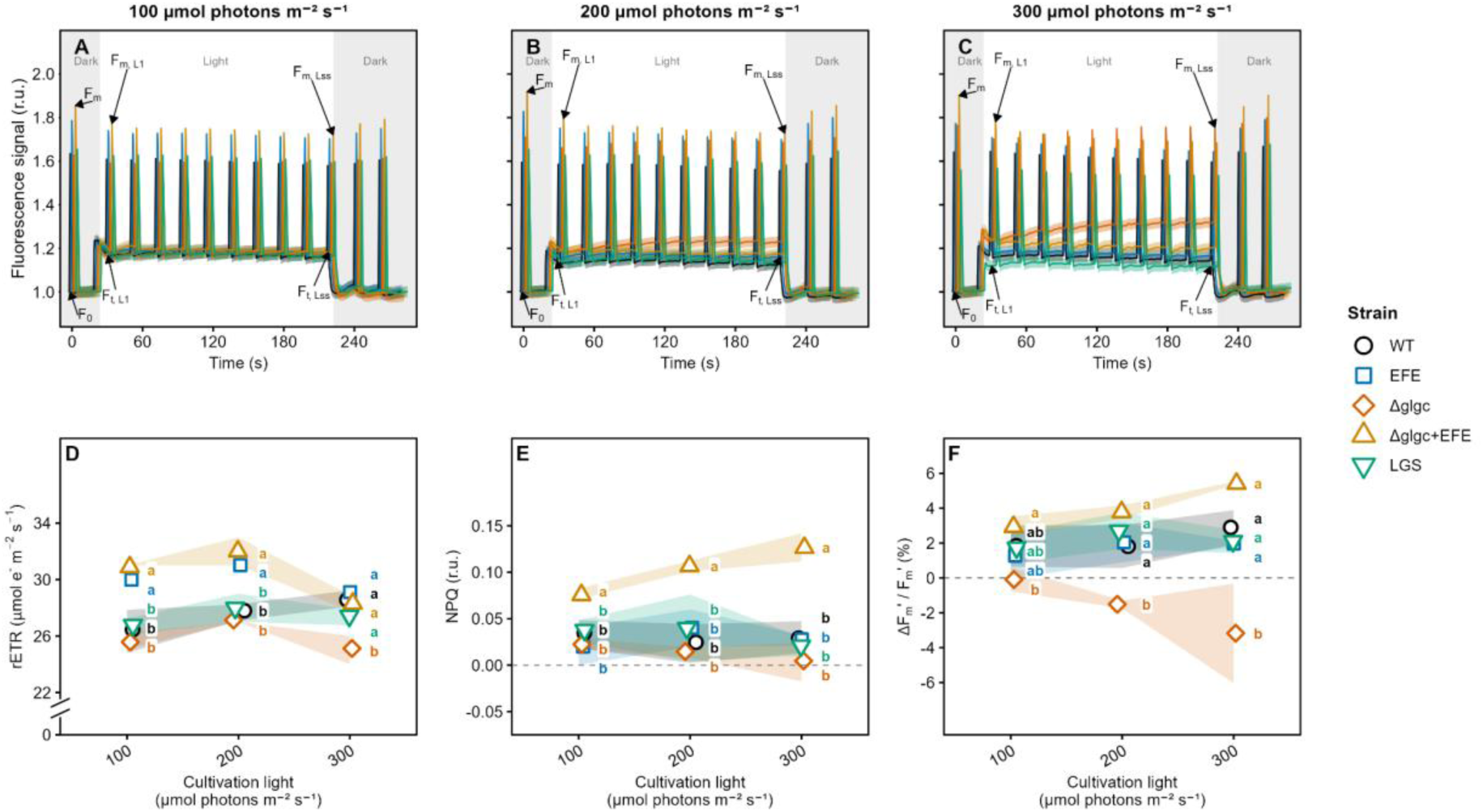
Slow (Kautsky) chlorophyll-fluorescence induction kinetics in the tested *Synechocystis* strains cultivated at 100-300 µmol photons m^-2^ s^-1^ (A-C), and derived parameters (D-F). Relative electron transport rate (rETR; panel D) and non-photochemical quenching (NPQ; panel E) were calculated from maximum and steady-state fluorescence values measured after light acclimation together with initial fluorescence (F_O_), according to Eqs. 1 and 2. The change in maximal fluorescence after light acclimation (ΔF_m_′/F_m_′; Eq. 4, panel F) was used as a proxy for the extent of state transitions (Calzadilla and Kirilovsky, 2020; Kis et al., 2026). Letters in panels D-F indicate significant differences between strains within each light intensity and n.s. (not significant) otherwise; n = 3. Further parameters are provided in Supplementary Figure S1.

Rapid light curves were measured at PFD of 10 - 1 000 µmol photons m^-2^ s^-1^. Parameters derived from the LC3 measurements included the initial slope of the rETR-PFD relationship at low irradiance (α), which reflects the apparent quantum efficiency of photosynthesis; the slope at high irradiance (β), associated with photoinhibition; and the maximum measured relative electron transport rate (rETR_max_). Parameters α and β were obtained by direct fitting of rETR as a function of PFD (Ralph and Gademann, 2005). The following parameters were derived from OJIP fluorescence transients:

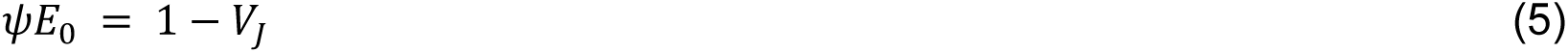

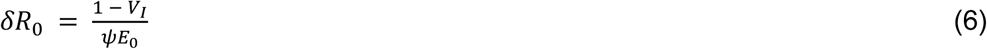

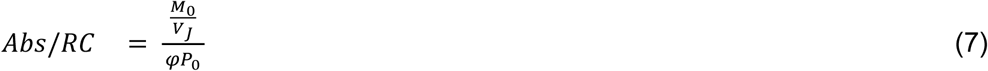

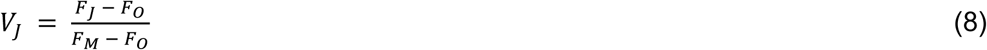

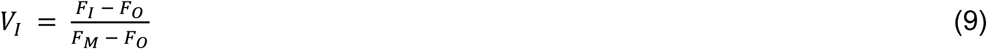

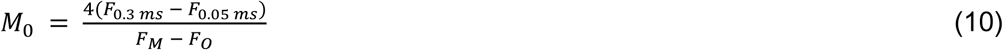

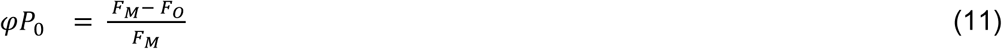

where ψE_0_ represents the efficiency of electron transfer from the primary quinone acceptor Q_A_^⁻^ to the plastoquinone pool, δR_0_ represents the efficiency of electron transfer from plastoquinol (PQH_2_) to the final electron acceptors of PSI, and Abs/RC represents the apparent antenna size of an active PSII reaction centre. The F_J_ and F_I_ denote fluorescence levels at the J and I steps of the OJIP transient, measured at 2 ms and 30 ms, respectively.

### 2.4 Ethylene production and cellular composition

Ethylene production by *EFE* and *ΔglgC+EFE* strains was measured by gas chromatography with a thermal conductivity detector (GC-TCD) using a Porapak N column and helium carrier gas, isothermal elution at 60 °C (Agilent 7890A). For ethylene measurements, 5 mL of culture was transferred to sealed 25 mL glass Hungate tubes and incubated overnight in the Percival cultivation chamber under the same conditions as the corresponding culture. For GC analysis, 500 μL of headspace gas was injected, and ethylene concentration was quantified using the retention time and peak area obtained from ethylene standards.

Chlorophyll *a* and total carotenoid contents were determined according to the previously described protocol (Zavřel et al., 2015). Storage polysaccharide content was determined in the cellular pellets remaining after pigment extraction, using the phenol–sulfuric acid method described previously (Zavřel et al., 2018a). Cell count was performed using a Cellometer Auto X4 automated cell counter (Nexcelom Bioscience, Lawrence, MA, USA), and cellular dry weight was determined gravimetrically from lyophilized biomass. Cell size was determined using a Multisizer 4e Coulter Counter (Beckman Coulter), with samples fixed in 2% formaldehyde. Lyophilized biomass was further analysed by Fourier-transform infrared spectroscopy (FTIR) using Nicolet IS10 spectrometer (Thermo Fisher Scien-tific) following previously described protocols (Felcmanová et al., 2017; Rabouille et al., 2021). Briefly, the amplitudes of FTIR bands at approximately 1735 cm^-1^, 1652 cm^-1^, and 1152 cm^-1^ were used as proxies for lipid, protein, and carbohydrate content, respectively. As protein content showed the lowest dependence on the experimental conditions among the three macromolecular pools (data not shown), spectra were normalized to the protein band at approximately 1652 cm^-1^. The relative fatty-acid chain length was estimated from the ratio of the amplitudes of the asymmetric CH_2_ and CH_3_ stretching vibrations at approximately 2925 cm^-1^ and 2960 cm^-1^, respectively. The relative content of *cis* unsaturated bonds was estimated from the amplitude of the olefinic =C–H stretching band at approximately 3010 cm^-1^, with higher values indicating a higher degree of acyl-chain unsaturation.

### 2.5 Proteomics

Proteomic analysis was performed using lyophilized biomass following the previously described protocol (Berková et al., 2026). Briefly, approximately 10 mg of dry biomass was washed with a methyl *tert*-butyl ether/methanol mixture (3:1, v/v) and extracted with 0.6 mL of SDS buffer containing 2% (w/v) SDS, 30% (w/v) sucrose, 5% (v/v) β-mercaptoethanol, 5 mM EDTA, and 100 mM Tris-HCl (pH 8.0), together with 0.3 mL of buffer-saturated phenol. Proteins in the phenolic phase were precipitated overnight at -20 °C using 1.6 mL of ice-cold 100 mM ammonium acetate in methanol. Pellets were washed with 0.5 mL of 80% (v/v) aqueous acetone, dried, and dissolved in a buffer containing 100 mM NH_4_HCO_3_, 8 M urea, and 10 mM DTT. Aliquots corresponding to 10 μg of protein were alkylated with iodoacetamide, diluted to a final urea concentration of 2 M, and digested with trypsin. The resulting peptides were loaded onto EvoTips and analysed using the Evosep One system with the 30 samples-per-day (SPD) method and an EV1137 Performance Column (Evosep Biosystems, Odense, Denmark), coupled to an Orbitrap Fusion Lumos Tribrid Mass Spectrometer equipped with a FAIMS Pro interface (Thermo Fisher Scientific, Waltham, MA, USA). Samples were analysed using alternating FAIMS compensation voltages of -40 and - 55 V.

The acquired MS/MS spectra were recalibrated and searched against the UniProt reference protein database (UP000001425_2026_03_14) and a list of common protein contaminants using Proteome Discoverer 2.5 and algorithms Sequest HT, MS Amanda 2.0 or MSFragger (Dorfer et al., 2014; Kong et al., 2017). Only proteins with at least two unique peptides (or one unique peptide and at least 10 peptides measured) were considered for quantitative analysis. Quantitative differences between proteins were determined using Minora precursor-ion quantification followed by normalization. The mass spectrometry proteomics data have been deposited in the ProteomeXchange Consortium via the PRIDE partner repository (Perez-Riverol et al., 2022) under the dataset identifier PXD076169.

### 2.6 Metabolic overflow measurement

To measure the metabolic overflow, cultivation medium from cultures growing at PFD 100 - 300 µmol photons m^-2^ s^-1^ was analyzed for the presence of organic compounds. Samples were spiked with ribitol as an internal standard to a final concentration of 0.1 ng µL^-1^, and 50 µL aliquots were dried in a speed-vacuum concentrator. Samples were derivatized as described previously (Dufková et al., 2023) and transferred to glass vials equipped with 300 µL glass inserts. A semi-targeted GC-MS workflow was used to profile tricarboxylic acid (TCA) cycle and energy-metabolism intermediates. Derivatized samples (0.4 µL) were analysed using a Zebron ZB-5MSplus GC column (Phenomenex; 30 m × 0.25 mm × 0.25 µm) with a 10 m guard column. Samples were injected in split mode (5:1) at 250 °C, using helium as the carrier gas at 1.2 mL min^-1^. The temperature program was: 70 °C for 5 min, 10 °C min⁻¹ to 220 °C, 40 °C min⁻¹ to 320 °C, and 320 °C for 5 min. Analytes were ionized via EI (70 eV, 50 µA). Data was processed using Compound Discoverer 3.3 (Thermo Fisher Scientific) with a mass tolerance of 5 ppm, a total ion chromatogram (TIC) threshold of 25 000, and a signal-to-noise (S/N) threshold of 3. TCA-cycle and energy-metabolism intermediates were identified against the NIST 2023, GC-Orbitrap, and in-house spectral libraries using a spectral match score >90 and a ΔRI <0.5%. Peak assignments were validated in Skyline 19.1 (Pino et al., 2020) using extracted-ion chromatograms with a mass tolerance of 5 ppm. Glucose was excluded from the analysis because of co-elution with components of the cultivation medium. Compounds whose mass spectra were consistent with carbohydrates but could not be confidently assigned to a specific structure were reported collectively as “carbohydrates”, as the electron-ionization spectra of derivatized carbohydrates are often highly similar across isomers, and unambiguous structural identification would require authentic reference standards or an orthogonal analytical method such as NMR.

### 2.7 Flux Balance Analysis of *Synechocystis* metabolism

Flux balance analysis (FBA) was based on the previously published genome-scale metabolic reconstruction of *Synechocystis* metabolism (Höper et al., 2024). The reconstruction was extended by adding the metabolites ethylene (C_2_H_4_) and guanidine (CH_6_N ^+^). In addition, the endogenous gene *sll1077* (*gdmH*), guanidine hydrolase, and the corresponding reaction converting guanidine to urea and ammonium were introduced. The heterologous *efe* gene encoding the ethylene-forming enzyme from *Pseudomonas syringae* pv *phaseolicola* PK2 was also added. The updated network was used for FBA to estimate the relative metabolic costs of individual metabolites. For each metabolite, a demand reaction was introduced and constrained to a flux of 1 mmol gDW^-1^ h^-1^. The model was then optimized to minimize photon uptake, and the resulting fluxes were extracted for comparison.

For comparison of the WT, *ΔglgC*, and *LGS* strains, glycogen was removed from the biomass reaction and represented by a separate boundary reaction while remaining included in the calculation of cellular mass. All simulations corresponded to cultivation at 200 µmol photons m⁻^2^ s⁻¹, for which the photon uptake rate was determined as the value reproducing the measured WT growth rate. The glycogen content of WT cells was calculated according to previous work (Höper et al., 2024), and the growth rate was constrained to the experimentally measured value of 0.049 h⁻¹. The model was first optimized to minimize photon uptake, and the resulting ATP production by ATP synthase and minimum NADPH production associated with ferredoxin:NADP⁺ oxidoreductase (FNR) were extracted. Glycogen production was subsequently constrained to zero for *ΔglgC* or to 50% of the WT value for *LGS*. A demand reaction for the metabolite of interest was then introduced, and its export flux was constrained to the value at which the ATP/NADPH ratio matched that of the WT model. The same procedure was applied to the *ΔglgC+EFE* strain, except that ethylene production was additionally constrained to the experimentally measured value before calculating the corresponding metabolite export flux.

### 2.8 Statistical analysis

All statistical analyses were performed in R v4.5.2. Unless otherwise stated, the significance threshold was set to α = 0.05. The main experimental dataset was analysed as a two-factor design with strain and cultivation light intensity as fixed factors. Multi-point instrument recording data sets (chlorophyll fluorescence recordings: LC3, NPQ3, OJIP) consisted of several individual measurements per replicate and directly derived parameters were thus treated as approximately normally distributed (Figures 2-4). For these parameters two-way ANOVA was used first, and significance values used for generating the compact letter displays were obtained by one-way ANOVA with Tukey’s HSD as a post-hoc test separately for each light intensity. The remaining variables were treated as non-normally distributed and were analysed by aligned rank transform (ART) two-way ANOVA (Wobbrock et al., 2011) using the ARTool package (Kay et al., 2025) with pairwise contrast being obtained by art.con() + Holm correction (Elkin et al., 2021).

**Fig. 3.**
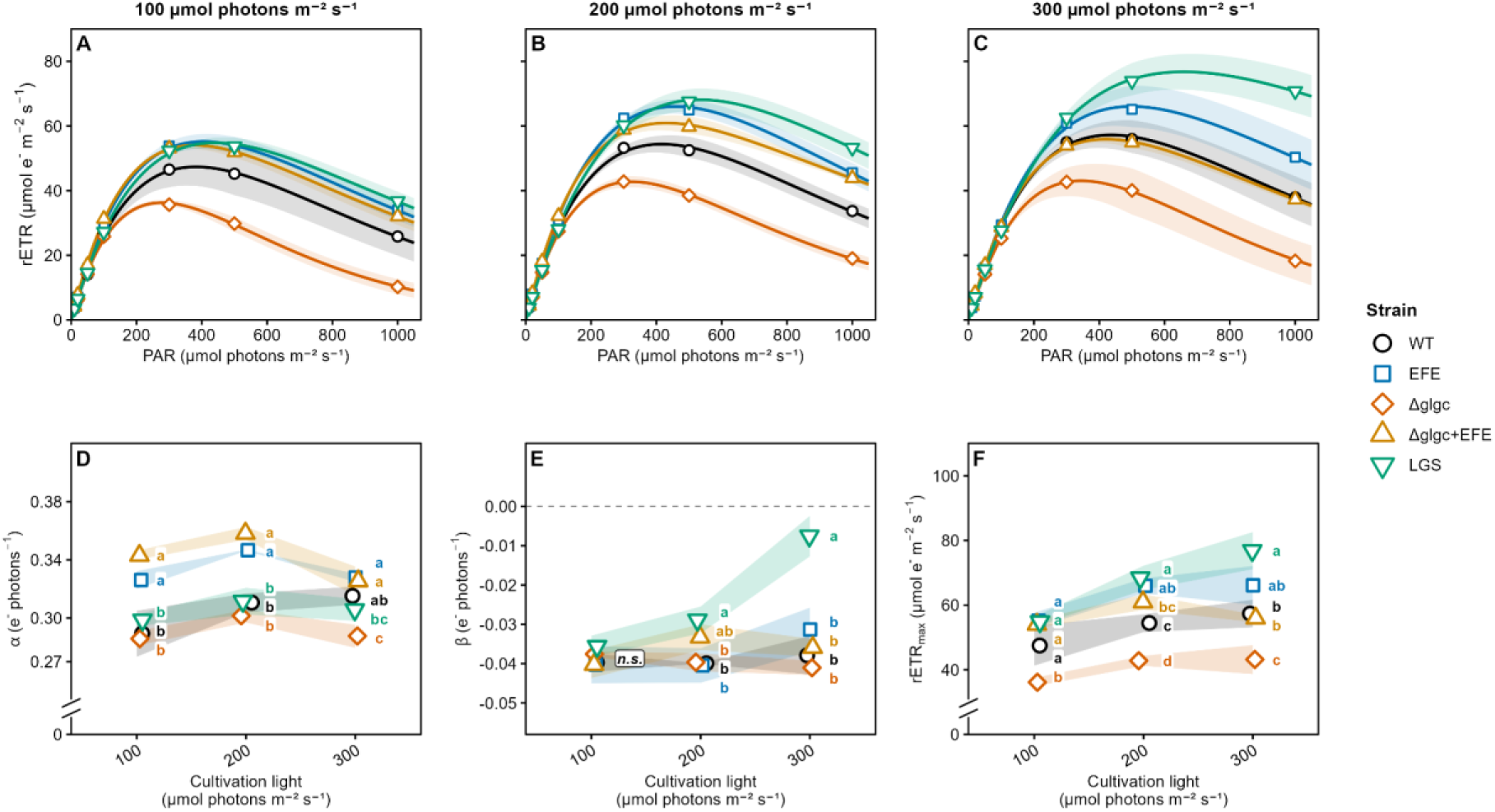
Relative electron transport rate in the tested *Synechocystis* strains as a function of irradiance measured using rapid light curves (A-C), and parameters derived from the curves for cultures cultivated at 100- 300 µmol photons m^-2^ s^-1^ (D-F). α, initial slope of the light curve and indicator of apparent quantum efficiency; β, slope at high irradiance and indicator of photoinhibition; rETR_max_, maximum relative electron transport rate ^(^_Ralp_^h^and Gademann, 2005). Letters in panels D-F indicate significant differences between strains within each light intensity and n.s. (not significant) otherwise; n = 3.

**Fig. 4.**
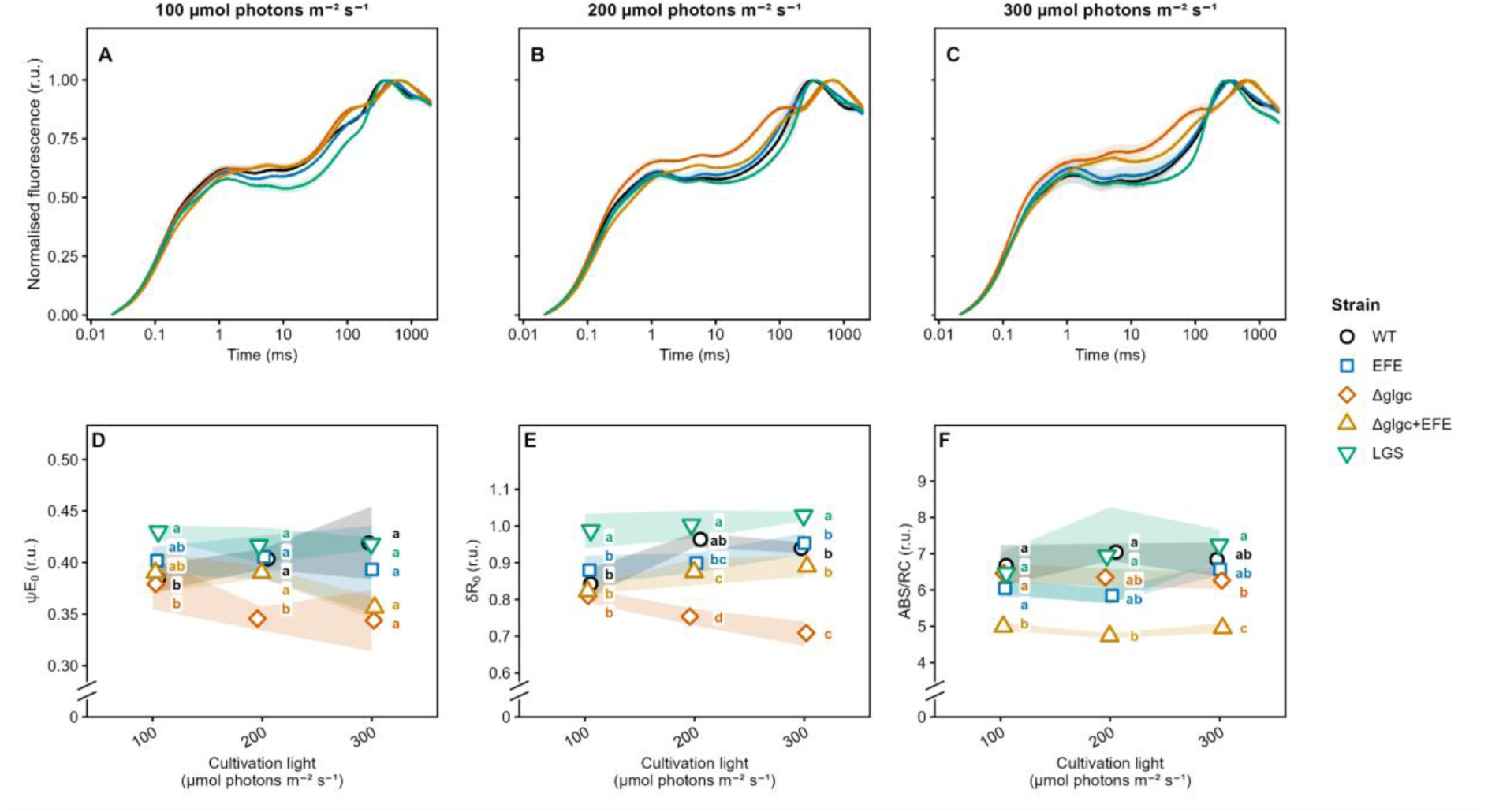
Fast fluorescence-induction kinetics (OJIP transients) in the tested *Synechocystis* strains cultivated at 100- 300 µmol photons m^-2^ s^-1^ (A–C) and derived JIP-test parameters (D–F). ψE_0_ and δR_0_ represent the efficiencies of electron transfer from Q_A_⁻ to the plastoquinone pool and from plastoquinol (PQH_2_) to the final PSI acceptors, respectively, while Abs/RC represents the apparent antenna size of active PSII (Tsimilli-Michael, 2020). Letters in panels D-F indicate significant differences between strains within each light intensity and n.s. (not significant) otherwise; n = 3. Further parameters are provided in Supplementary Figure S2.

## 3. Results

### 3.1 Glycogen downregulation increases growth rate under low light but impairs growth under high light

Strains were cultivated across a PFD range of 35-300 µmol photons m^-2^ s^-1^. At low irradiance (35 and 100 µmol photons m^-2^ s^-1^) both *ΔglgC* and *LGS* strains exhibited slightly higher specific growth rates than WT (0.022±0.003 h^-1^ and 0.020±0.002 h^-1^ vs. 0.018±0.003 h^-1^, respectively; Fig. 1). Above 100 µmol photons m^-2^ s^-1^, WT growth continued to increase, reaching 0.054±0.003 h^-1^ at 300 µmol photons m^-2^ s^-1^, whereas *ΔglgC* growth declined sharply to 0.020±0.003 h^-1^ at the highest tested irradiance. In contrast, *LGS* showed only a moderate reduction at 300 µmol photons m^-2^ s^-1^: 0.044±0.010 h^-1^ (Fig. 1A). This light dependence is consistent with previous reports showing that glycogen-deficient cyanobacteria increasingly redirect fixed carbon into overflow metabolism as photon supply rises. In *Synechocystis* and *Synechococcus* sp. PCC 7002, *glgC* deletion increased metabolic overflow, with increasing irradiance promoting excretion of 2-OG and pyruvate (Gründel et al., 2012; Jackson et al., 2015); in *Synechocystis*, increasing photon supply from 50 to 250 µmol photons m^-2^ s^-1^ was sufficient to induce extracellular accumulation of these metabolites (Cano et al., 2018). As the excreted organic acids represent fixed carbon lost from biomass, together with the ATP and NADPH invested in fixing it, this progressive diversion provides a plausible basis for the growth penalty observed here at elevated irradiance. In the present experiment, the transition from a growth advantage to a growth penalty in *ΔglgC* occurred between 100 and 200 µmol photons m^-2^ s^-1^. Notably, the light–growth responses of *ΔglgC* and *ΔglgC+EFE* were not captured well by conventional saturation–photoinhibition relationships such as Aiba/Haldane (Andrews, 1968). Instead, both strains exhibited a threshold-like response, with growth declining abruptly above a critical irradiance rather than gradually entering photoinhibition (Fig. 1A). The Platt model (Platt et al., 1980) was nevertheless used to obtain comparable light-response parameters across strains, while recognising that its photoinhibition term does not fully describe the sharp suppression observed in the glycogen-deficient backgrounds.

The EFE strain grew consistently slower than WT across the irradiance range, with growth reduced by approximately 10–18%, indicating a substantial metabolic burden associated with EFE expression (Zavřel et al., 2016a; Sebesta et al., 2024). The combined *ΔglgC+EFE* strain showed the strongest growth impairment at 100-300 µmol photons m^-2^ s^-1^. At the lowest irradiance, however, it grew slightly faster than EFE alone and at a rate comparable to WT (Fig. 1A). Thus, under low light, reducing glycogen synthesis partially offset the growth burden imposed by EFE expression, whereas at elevated irradiance, the effects of the two interventions became additive.

Storage polysaccharide content, determined at 200 µmol photons m^-2^ s^-1^, was approximately 575 mg gDW^-1^ in WT and *EFE* and approximately 60–80 mg gDW^-1^ in *ΔglgC*, *ΔglgC+EFE* and *LGS* (Supplementary Figure S3), consistent with the intended glycogen phenotypes and with *LGS* not yet having restored glycogen synthesis at this irradiance. The residual signal in the glycogen-deficient strains is expected, as the method does not fully distinguish glycogen from other cellular polysaccharides. Replicate variability was high and the differences were not statistically significant, so these values are reported as a qualitative confirmation of the genotypes.

### 3.2 EFE expression in the *ΔglgC* background increases ethylene production but further compromises growth

Glycogen removal enhanced ethylene production most strongly at intermediate irradiance. At 200 µmol photons m⁻² s⁻¹, *ΔglgC+EFE* produced about 56% more ethylene than *EFE* (46±6 vs. 30±9 µmol(C_2_H_4_) gDW^-1^ h^-1^, respectively; Fig. 1B). Quantitative proteomics, performed at 200 and 300 µmol photons m⁻² s⁻¹, showed that this gain was achieved with equal or lower investment in the enzyme itself: EFE accounted for 0.8-1.2% of the total protein pool in *ΔglgC+EFE*, compared with 1.2-1.4% in *EFE* at 200 and 300 µmol photons m⁻² s⁻¹, respectively. Higher ethylene production in *ΔglgC+EFE* was therefore achieved without a corresponding increase in EFE abundance. Absolute rates are not directly comparable with earlier flat-panel photobioreactor measurements (70-260 µmol(C_2_H_4_) gDW^-1^ h^-1^ (Zavřel et al., 2016b)), which used a different construct and quantified ethylene in real time by membrane-inlet mass spectrometry rather than from headspace accumulation in sealed vials. Light quality may also contribute to the difference (Rodrigues et al., 2023; Zavřel et al., 2024).

The advantage of the *ΔglgC* background was irradiance-dependent. At 100 µmol photons m⁻² s⁻¹, the two strains performed equivalently (32±14 and 32±11 µmol C_2_H_4_ gDW^-1^ h^-1^ in *EFE* and *ΔglgC+EFE*, respectively; Fig. 1B), consistent with the benefit of glycogen removal emerging once photon supply exceeds the capacity of the native carbon sinks. At 300 µmol photons m⁻² s⁻¹, both strains produced more ethylene than at lower irradiance, and *ΔglgC+EFE* retained an advantage of approximately 16% (50±8 vs. 43±3 µmol(C_2_H_4_) gDW^-1^ h^-1^). Glycogen removal thus enhanced carbon partitioning toward ethylene across the upper part of the tested light range, with the largest relative gain at 200 µmol photons m⁻² s⁻¹. The gains in ethylene production were nevertheless smaller than the accompanying increase in photosynthetic electron transport in the EFE-containing strains (Figures 2-3), showing that part of the additional electron flow reached the product while the remainder was accommodated elsewhere.

This irradiance-dependent enhancement under nitrogen-replete conditions differs from earlier observations under nitrogen limitation, where ethylene production in *ΔglgC+EFE* was reduced despite elevated intracellular 2-OG (Veetil et al., 2017). Conversely, increased product formation in *ΔglgC* backgrounds under nitrogen limitation has been reported for other compounds, including lactate (van der Woude et al., 2014) and ethanol in a *ΔglgCΔphaCE* strain (Namakoshi et al., 2016). These contrasting outcomes indicate that the effect of glycogen removal on product formation depends strongly on the product pathway and on the physiological context in which the strain is operated.

Taken together, the results show that under nitrogen-replete conditions glycogen removal enhances carbon partitioning toward ethylene in the EFE background at elevated irradiance, most effectively at 200 µmol photons m⁻² s⁻¹. EFE can therefore provide an additional sink for the ATP, NADPH, and carbon made available by reducing glycogen synthesis. However, this sink is insufficient to accommodate the entire surplus. Consequently, additional metabolic sinks must contribute to balancing the excess carbon and reducing equivalents, or overall carbon fixation must decline, both of which are consistent with the reduced growth observed in *ΔglgC+EFE* at elevated irradiance.

### 3.3 EFE expression stimulates photosynthetic electron transport by redistributing electron flux rather than remodelling the photosynthetic apparatus

Both EFE-containing strains exhibited increased relative electron transport rates (rETR; Eq. 1) under actinic illumination (Fig. 2A-D). They also showed higher α values (Fig. 3A-D) , reflecting increased initial slopes of the rapid light curves and therefore higher apparent quantum efficiency of photosynthesis (Ralph and Gademann, 2005). At 100-200 µmol photons m^-2^ s^-1^, rETR and α were approximately 10–15% and 11–20% higher, respectively, in EFE and *ΔglgC+EFE* than in the other strains. Both parameters declined at the highest irradiance but remained above those of the other strains. In contrast, *ΔglgC* consistently showed the lowest values. rETR was comparable in the two EFE-containing strains across the tested irradiances, although their ethylene production rates differed by up to 56% (Fig. 1B), placing the determinant of product formation downstream of electron supply. These differences in rETR and α were not apparent in the OJIP transients, which were measured after dark acclimation (Fig. 4).

The increased rETR and α in *EFE* and *ΔglgC+EFE* strains were not accompanied by detectable changes in the apparent size of active PSII (Fig. 4F) or in the abundance of proteins associated with light harvesting and the photosynthetic electron transport (PSET) chain (Fig. 5B). The increase in electron transport is therefore consistent with the carbon-sink effect (Grund et al., 2022), where introduction of a heterologous NADPH-consuming pathway increases photosynthetic electron transport capacity. This agrees with the broader observation that cyanobacterial photosynthesis often operates below its maximal capacity and can be stimulated by increased downstream electron demand, particularly under light-limiting conditions (Grund et al., 2019). The absence of substantial changes in antenna size or PSET- protein abundance suggests that the elevated rETR and α primarily reflect redistribution of electron flux rather than major structural remodelling of the photosynthetic apparatus, consistent with the flexibility of cyanobacteria metabolism (Xiong et al., 2017).

**Fig. 5.**
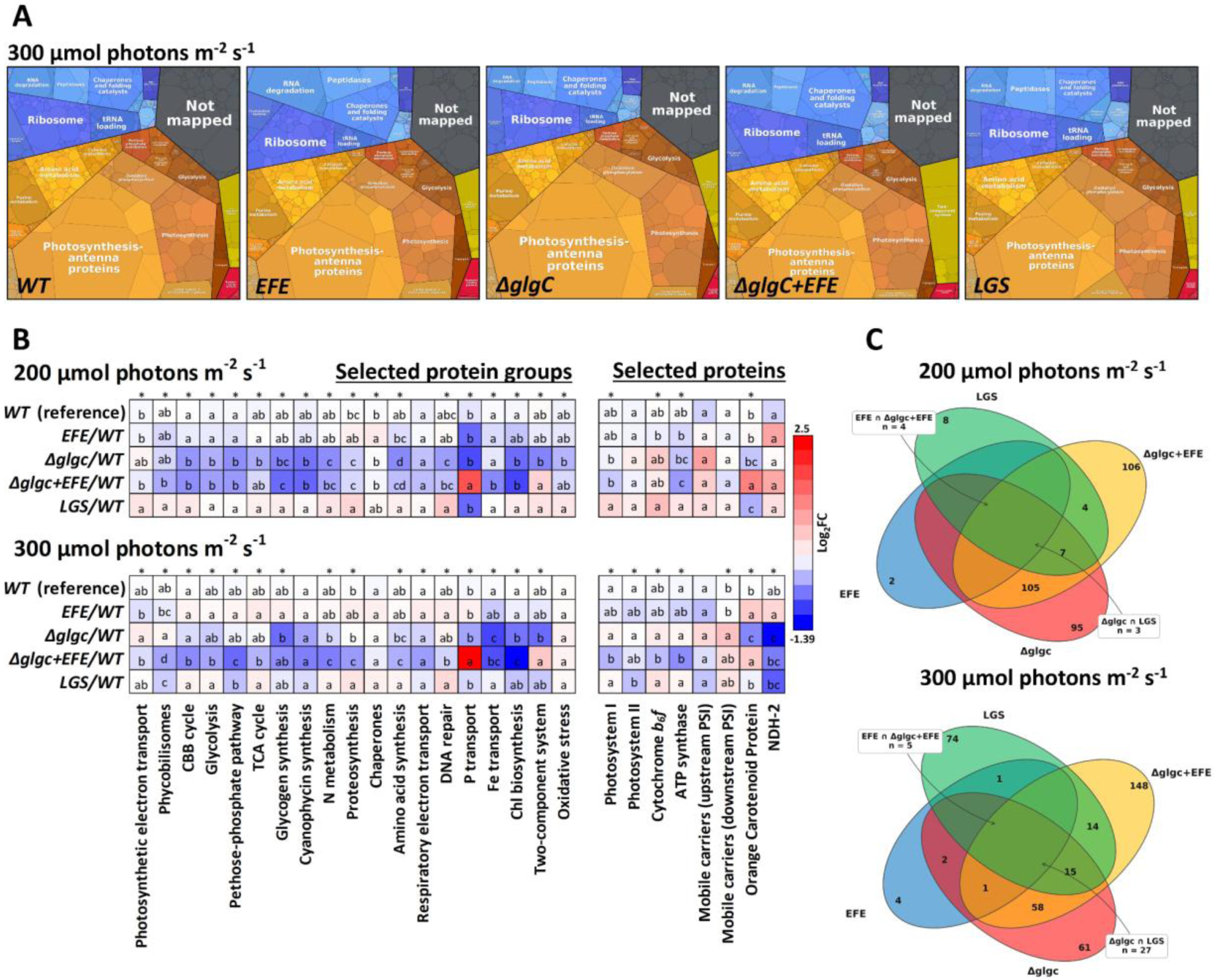
Proteomic analysis of the tested *Synechocystis* strains. The relative distribution of quantified proteins according to KEGG annotation was visualized using Proteomaps (Liebermeister et al., 2014) (A). Proteins were further grouped into 25 predefined functional categories (B; Supplementary Table S1); category-level log2-FC relative to WT was calculated for each biological replicate as log2(replicate intensity / mean WT intensity) and summarized as the median log₂FC across all proteins within each category (four replicates per strain; 438 of 2 368 detected proteins assigned; 200 and 300 µmol photons m-2 s-1 conditions normalised independently). WT-noise values represent within-WT replicate variability and provide a baseline reference. Letters indicate ANOVA post-hoc groupings for individual categories; asterisks indicate overall significance (*p* < 0.05). Overlap of significantly changed proteins between strains (|log2-FC| ≥ 1.0, BH-adjusted *p* < 0.05) at each light intensity is shown as Venn diagrams (C). Numbers indicate proteins shared exclusively by the indicated combinations of strains or unique to individual strains. The two diagonal overlaps in the four-set diagram represent EFE ∩ *ΔglgC+EFE* and *ΔglgC* ∩ *LGS*. The top 50 proteins showing the largest changes between WT and the tested strains are provided in Supplementary Figure S4; complete volcano plots are provided in Supplementary Figure S5.

### 3.4 EFE expression in the *ΔglgC* background induces secondary phosphate limitation, activating photoprotection and reducing potential ethylene production

NPQ in *ΔglgC+EFE* increased steeply with irradiance, from below the level of *EFE* at 200 µmol photons m^-2^ s^-1^ to the highest of all strains at 300 µmol photons m^-2^ s^-1^ (Fig. 2E). Between these two irradiances the ethylene gain over *EFE* fell from about 56% to 16% (Fig. 1B), indicating a shift from product formation toward dissipation as irradiance increased. This contrasted with *ΔglgC*, which exhibited low NPQ across the tested light range. In cyanobacteria, NPQ is mediated primarily by the orange carotenoid protein (OCP) at the level of the phycobilisome (PBS) (Domínguez-Martín et al., 2022), and is induced by blue-green light and high irradiance (Kirilovsky and Kerfeld, 2016). Consistent with the fluorescence results, OCP was most strongly upregulated relative to WT in *ΔglgC+EFE* (Fig. 5B). OCP abundance also increased in EFE, but to a lesser extent, whereas *ΔglgC* showed progressive OCP downregulation with increasing irradiance.

The *ΔglgC+EFE* strain also displayed the strongest induction of phosphate-transport proteins, including all components of the Pst1 and Pst2 systems (Fig. 5B; Supplementary Table S1). This proteomic pattern is consistent with intracellular inorganic phosphate (Pi) limitation. We propose that continuous EFE-mediated 2-OG consumption, together with extracellular 2-OG export, competes with nitrogen assimilation for substrate through the GS-GOGAT cycle (Veetil et al., 2017). Reduced GS-GOGAT flux can be expected to limit glutamine availability for nucleotide synthesis and amino-acid production, while high-light stress increases the demand for biosynthetic repair. These processes can be expected to increase cellular dependence on phosphate recycling and thereby contribute to depletion of the free Pi pool. Consistent with this interpretation, exopolyphosphatase Ppx was strongly induced (Supplementary Table S1), indicating mobilisation of polyphosphate as a potential emergency Pi source (Sanz-Luque et al., 2020), whereas polyphosphate kinase (PPK) remained close to WT levels (Supplementary Table S1). As protein abundance does not report flux, these data do not distinguish net polyphosphate degradation from continued cycling between synthesis and degradation.

The *ΔglgC+EFE* strain also showed the largest reduction in apparent PSII antenna size (Abs/RC; Fig. 4F), likely associated with broader downregulation of PBS proteins (Fig. 5B). At the same time, it exhibited the strongest State 2→1 transition under actinic illumination (Fig. 2F). This suggests that *ΔglgC+EFE* attempted to maintain functional coupling of the remaining PBS antenna to PSII while reducing overall excitation pressure. Together with the strong OCP-mediated NPQ response, these changes indicate that *ΔglgC+EFE* relies on multiple layers of photoprotection under high irradiance.

Thus, EFE expression in the *ΔglgC* background has two contrasting effects. It provides an additional sink for carbon and reducing equivalents and increases ethylene production, but the associated 2-OG drain is accompanied by a phosphate-stress response, enhanced photoprotection, and reduced investment in the photosynthetic apparatus. These responses provide a plausible explanation for the reduced growth and incomplete conversion of additional photosynthetic electron flow into ethylene at elevated irradiance.

### 3.5 *ΔglgC* strain controls PSII flux through state transitions and reduces investment in iron-demanding machinery

The *ΔglgC* strain showed reduced rETR and α, particularly at the highest irradiance (Figures 2- 3), together with low NPQ and progressive OCP downregulation (Figures 2 and 5). The low NPQ was unexpected because removal of the glycogen sink could increase excitation pressure and oxidative stress at PSII, potentially stimulating photoprotective quenching (Pospíšil, 2012). Instead, *ΔglgC* was the only strain showing a clear State 1→2 transition under actinic illumination, corresponding to reduced PBS attachment to PSII (Fig. 2F). This was accompanied by the lowest qP values across the tested strains (Supplementary Figure S1), indicating a larger fraction of closed PSII centres. As the abundances of PBS and PSET proteins remained close to WT (Fig. 5B), the state transitions represented the primary mechanism for limiting PSII excitation in *ΔglgC* without major structural remodelling of the light-harvesting apparatus.

This interpretation agrees with previous observations that dark-acclimated *ΔglgC* cells retain PSII electron flow similar to WT but show reduced PSII quantum yield under actinic illumination (Cano et al., 2018). In the present study, *ΔglgC* also exhibited the highest V_J_ values after dark acclimation (Supplementary Figure S2), indicating a more reduced plastoquinone pool (Tóth et al., 2007). The low ψE_0_ and δR_0_ values indicated reduced electron-transfer efficiency from PSII to the plastoquinone pool and from plastoquinol to the final PSI acceptors, respectively (Fig. 4D-E; Eq. 5-6). The combination of a high V_I_ value and a weak IP phase (Fig. 4A-C) is consistent with limitation of electron flow at the PSI acceptor side (Schansker et al., 2005). Because FNR and Calvin–Benson–Bassham-cycle (CBB) enzymes become inactive during dark acclimation in *Synechocys*tis (Zavřel et al., 2026), the elevated V_I_ may reflect reduced cyclic electron flow (CEF) in *ΔglgC*, consistent with previous observations (Cano et al., 2018).

The proteomic data provide additional support for reduced CEF. Type II NADH oxidoreductase (NDH-2), which contributes to CEF in *Synechocystis* (Nikkanen et al., 2021), was downregulated in *ΔglgC*, and several NDH-1 subunits were likewise reduced (Fig. 5B, Supplementary Table S1). Whether respiratory electron transport (RET) also increased to drain plastoquinol remains less clear. Cytochrome *c* oxidase was slightly reduced, cytochrome *bd* oxidase was unchanged, and no other respiratory-chain components showed consistent upregulation (Fig. 5B, Supplementary Table S1). Although RET can increase without corresponding protein accumulation because terminal oxidases may operate below maximal capacity (Lea-Smith et al., 2013), the absence of increased abundance of terminal oxidases argues against a major increase in respiratory capacity. We therefore favour a model in which state transitions and reduced CEF limit PSII electron input, while overflow metabolism provides a downstream sink for carbon and reducing equivalents that are no longer committed to glycogen.

At the highest irradiance, *ΔglgC* also showed coordinated downregulation of iron-demanding proteins (Fig. 5B, Supplementary Table S1). This response is consistent with reduced cellular demand for iron-containing respiratory and biosynthetic machinery (Cano et al., 2018). The strain showed reduced abundance of respiratory components and proteins involved in light-independent chlorophyll biosynthesis through the protochlorophyllide reductase complex (Bröcker et al., 2010; Borisov et al., 2011; Schuller et al., 2019), accompanied by reduced abundance of iron-acquisition proteins. This pattern differs from classical iron limitation, in which IsiA induction and PSI-antenna remodelling are prominent (Bibby et al., 2001); neither response was observed here. We therefore interpret the reduction in iron-transport proteins as a demand-driven response rather than as evidence of iron starvation. Notably, the core photosynthetic machinery required for CO_2_ fixation and overflow metabolism, including iron-rich PSI, was maintained (Gao, 2020). This coordinated response was specific to *ΔglgC* and largely absent in *LGS*, consistent with the retention of partial glycogen flux in *LGS*.

### 3.6 Light-responsive attenuation of *glgC* in *LGS* provides a growth advantage under low irradiance but fails to maintain WT growth under high light

The *LGS* strain was designed to alleviate the fitness penalty of complete glycogen deletion by restoring *glgC* expression under control of the light-responsive *psbA2* promoter (Carrieri et al., 2012; Cantrell et al., 2023). At low irradiance (35-100 µmol photons m^-2^ s^-1^), this attenuation strategy combined reduced glycogen flux with sustained photosynthetic performance and resulted in higher growth than WT (Fig. 1A). *LGS* also exhibited the highest rETR_max_, the lowest β values, and the highest δR_0_ values among the strains tested (Figures 3-4), indicating efficient photosynthetic electron transfer and comparatively low photoinhibition even at high irradiance. Nevertheless, at 300 µmol photons m^-2^ s^-1^, LGS growth declined below the WT rate (Fig. 1A).

This decline coincided with light-dependent restoration of glycogen metabolism. Proteins associated with glycogen metabolism were reduced relative to WT at 200 µmol photons m^-2^ s^-1^ but converged toward WT levels at 300 µmol photons m^-2^ s^-1^ (Fig. 5B). This pattern is consistent with induction of the *psbA2* promoter at high irradiance (Mulo et al., 2009; Cantrell et al., 2023) and with partial restoration of glycogen synthesis (Cano et al., 2018; Cantrell et al., 2023). Restored glycogen flux would be expected to shift *LGS* toward WT phenotype at high irradiance; the fact that growth nevertheless remained lower than in WT suggests that additional metabolic responses contribute to the high-light phenotype.

At intermediate irradiance (100 and 200 µmol photons m^-2^ s^-1^) LGS accumulated more lipids and showed increased fatty-acid chain length and *cis* double bond content (Supplementary Figure S3), indicating that the accumulated lipid pool was also compositionally altered. Both processes consume reducing equivalents, with each desaturation step requiring reduced ferredoxin (Table 1). Enhanced lipid synthesis has previously been associated with compensation for reduced ATP and NADPH consumption under increased excitation pressure on PSII, including under CO_2_ limitation (Zavřel et al., 2018b). Considering the limited extracellular overflow metabolites detected in *LGS* (Fig. 6) and similar abundance of glycogen-metabolism proteins in *LGS* and WT (Fig. 5B), increased lipid synthesis represents a plausible intracellular sink for excess carbon and reducing equivalents at intermediate irradiance.

**Fig. 6.**
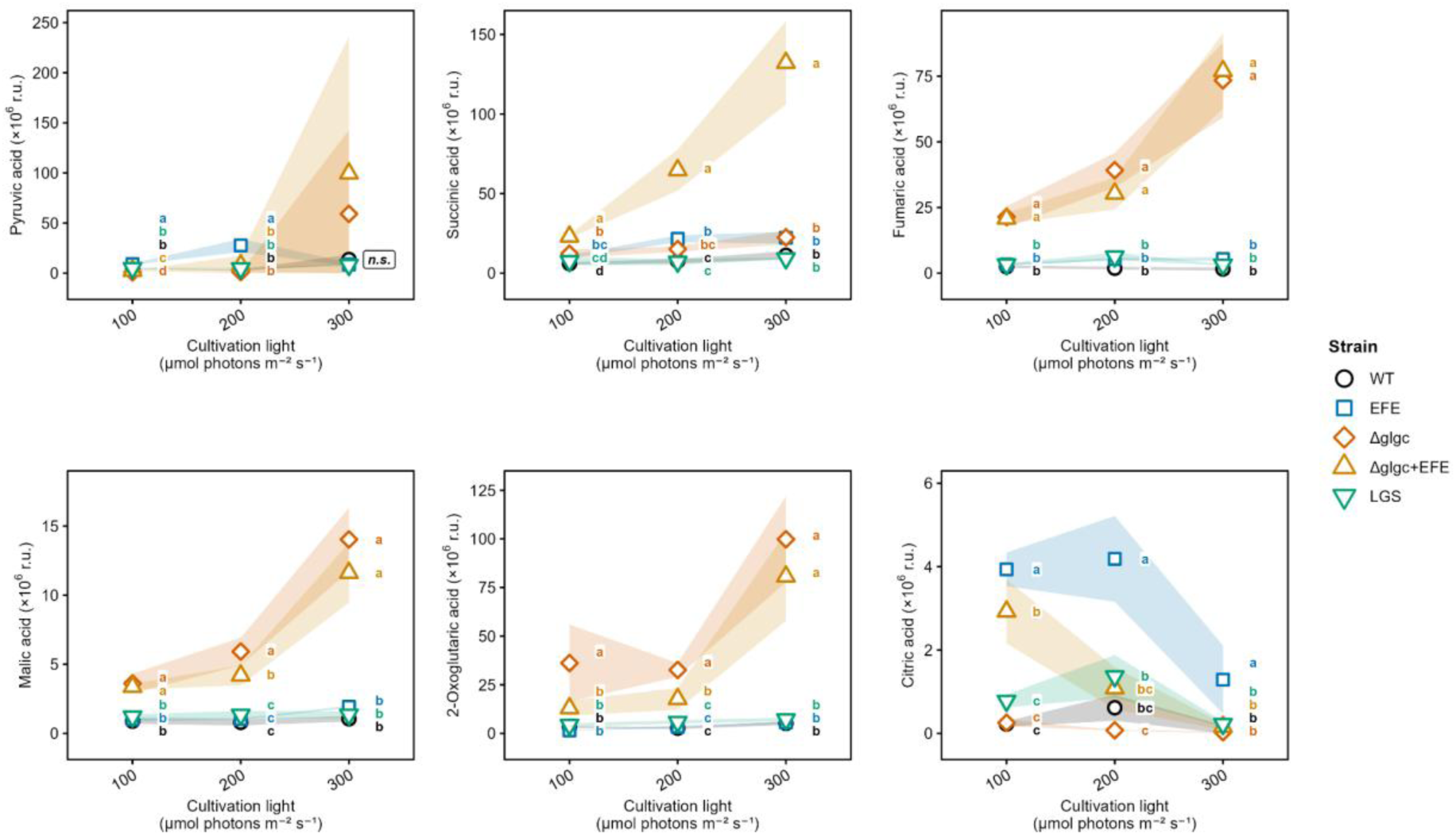
Overflow metabolites measured in the cultivation medium of the tested strains *Synechocystis* strains at 100- 300 µmol photons m^-2^ s^-1^. Values are expressed relative to ribitol as internal standard. Data represent averages ± SD; n = 4. Letters indicate significant differences between strains within each light intensity (p < 0.05); n.s., no significant pairwise differences at the indicated light intensity. In addition to the organic acids, also carbohydrate of unknown composition was detected in the cultivation medium (Supplementary Figure S6).

**Table 1.** Biosynthetic costs of overflow metabolites in the *Synechocystis* sp. PCC 6803 metabolic model compared with glycogen. Demand reactions represent ATP and NADPH requirements for synthesis of each metabolite from CO_2_, as described in the ‘Biosynthetic route’ column. Cost calculations are consistent with previous studies (Carrieri et al., 2015; Knoop and Steuer, 2015; Cano et al., 2018; Höper et al., 2024; Alford et al., 2025). Abbreviations: ACP, acyl carrier protein; ATP, adenosine triphosphate; EFE, ethylene-forming enzyme; F6P, fructose-6- phosphate; Fd, ferredoxin; G1P, glucose-1-phosphate; G3P, glyceraldehyde-3-phosphate; L-Arg, L-arginine; NADPH, nicotinamide adenine dinucleotide phosphate (reduced); OAA, oxaloacetate; PEP, phosphoenolpyruvate; 3-PGA, 3-phosphoglycerate; 2-OG, 2-oxoglutarate.

| Metabolite | Biosynthetic route from G3P | C atoms<br>(n) | ATP demand<br>(mol mol<sup>-1</sup>) | NADPH demand<br>(mol mol<sup>-1</sup>) | ATP/NADPH<br>(mol mol<sup>-1</sup>) | Photons<br>(mol mol<sup>-1</sup>) |
| --- | --- | --- | --- | --- | --- | --- |
| *Reference metabolite* |  |  |  |  |  |  |
| Glycogen | G3P → F6P → G1P → ADP-glucose → glycogen | 6 | 18.0 | 12.0 | 1.5 | 51 |
| *Major overflow metabolites* |  |  |  |  |  |  |
| Pyruvate | G3P → 3-PGA → PEP → pyruvate | 3 | 7.0 | 5.1 | 1.4 | 21 |
| Succinate | G3P → PEP → OAA → malate → fumarate → succinate | 4 | 14 | 7.2 | 1.9 | 34 |
| 2-Oxoglutarate | G3P → PEP → OAA; G3P → pyruvate → acetyl-CoA; OAA + acetyl-CoA → citrate → 2-OG | 5 | 14.0 | 8.1 | 1.7 | 37 |
| Fumarate | G3P → PEP → OAA → malate → fumarate | 4 | 8.0 | 6.2 | 1.3 | 25 |
| *Minor overflow metabolites* |  |  |  |  |  |  |
| Malate | G3P → PEP → OAA → malate | 4 | 8 | 6.2 | 1.3 | 25 |
| Citrate | G3P → PEP → OAA; G3P → pyruvate → acetyl-CoA; OAA + acetyl-CoA → citrate | 6 | 14 | 9.3 | 1.5 | 40 |
| *Macromolecular sinks* |  |  |  |  |  |  |
| Lipid (per saturated fatty acid, C16:0) | G3P → pyruvate → acetyl-CoA (× 8) → palmitoyl-CoA (C16:0) via fatty acid synthase | 16 | 55.0 | 46.0 | 1.2 | 184 |
| Lipid prolongation | Myristyl-ACP + Malonyl-ACP → palmitoyl-ACP | 2 | 7.0 | 6.0 | 1.2 | 24 |
| Each additional C=C double bond | fatty acyl chain + Fdred + O<sub>2</sub> → fatty acyl chain (+1 C=C) + Fdox + H<sub>2</sub>O | 0 | 1.3 | 1.0 | 1.3 | 4 |
| Protein (per average amino acid) | G3P → ... → amino acid (biosynthesis) | 5 | 18.2 | 8.8 | 2.1 | 64 |
| *Ethylene production (heterologous pathway)* |  |  |  |  |  |  |
| Ethylene | G3P → ... → 2-OG + L-Arg → C<sub>2</sub>H<sub>4</sub> (EFE) | 2 | 18.3 | 9.3 | 2.0 | 44 |

**Table 2.**
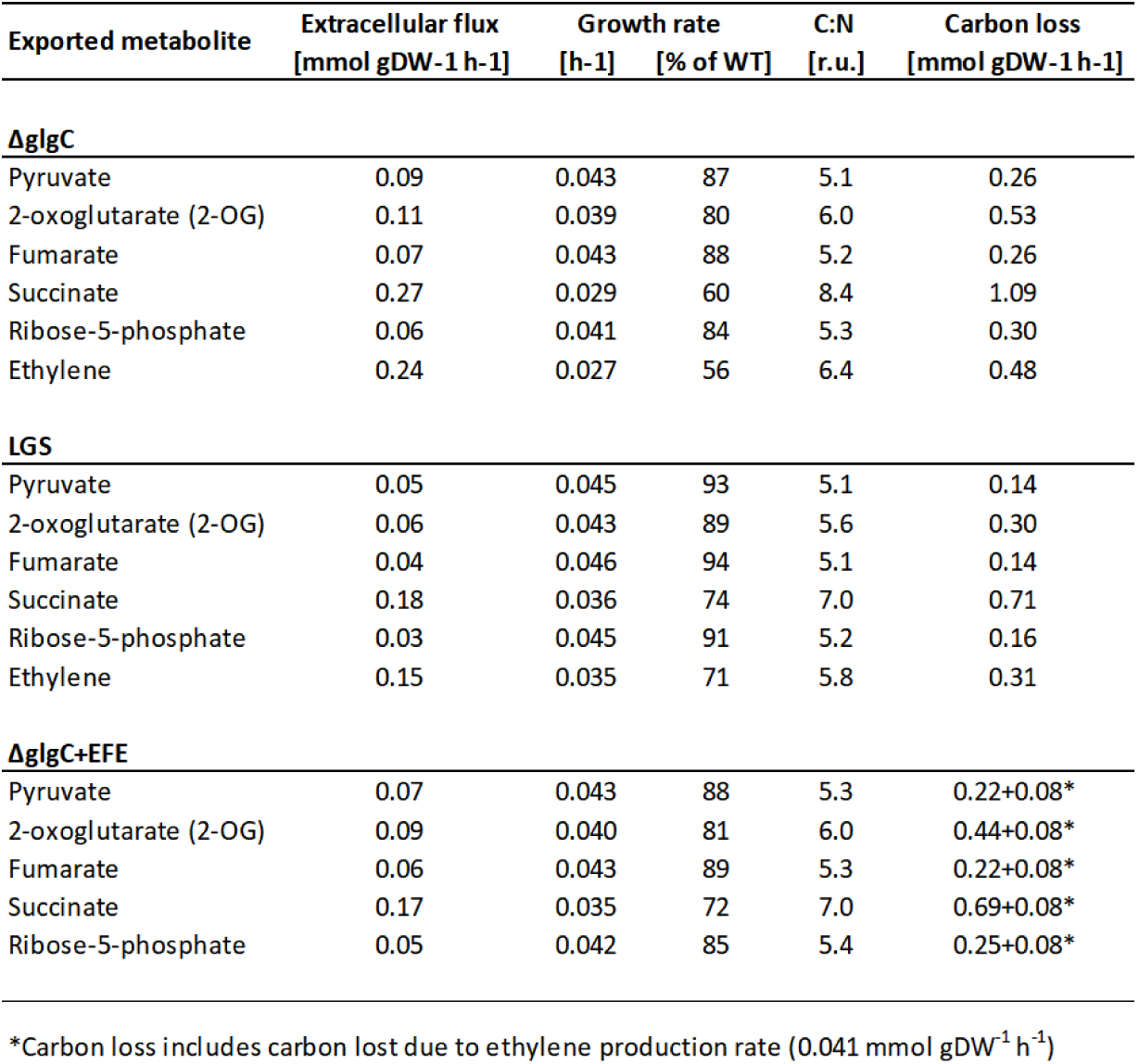
Simulated metabolite export required to restore the wild-type energy balance in strains with reduced or absent glycogen synthesis. Flux balance analysis was performed as described in Material & Methods - FBA. Each row represents an independent simulation in which the indicated metabolite acts as the sole energy and electron sink, and ethylene is included for *ΔglgC* and *LGS* as a hypothetical reference, although these strains do not carry *efe*. For *ΔglgC+EFE*, ethylene export was instead fixed at the measured rate (0.041 mmol gDW^-1^ h^-1^) before the export flux of each remaining metabolite was calculated, so ethylene does not appear as a separate row; the carbon released as ethylene (0.082 mmol C gDW^-1^ h^-1^, two carbons per molecule) is given as an additional term in the carbon-loss column. Ribose-5-phosphate serves as a proxy for excretion of pentose sugars. All fluxes are model predictions rather than measured values. “% of WT” indicates growth rate compared to the measured one for the indicated strain without excretion. r.u., relative units.

At 300 µmol photons m^-2^ s^-1^, both lipid content and lipid composition in *LGS* reached levels of the other strains. At the same time, *LGS* exhibited the highest cellular dry weight but not cell volume (Supplementary Figure S3). The concurrent restoration of glycogen-related protein abundance (Fig. 5B) and increase in the carbohydrate ratio are consistent with renewed investment in glycogen at high irradiance. Thus, carbon and reducing equivalents that may have been diverted into lipid synthesis at intermediate irradiance appear to be increasingly allocated to glycogen storage as *psbA2* activity increased.

Beyond the photosynthetic apparatus, *LGS* showed increased abundance of enolase (Eno) and the 2-OG/NtcA-responsive regulator Rre37 (Sll1330; Supplementary Table S1). Eno, which connects 2-phosphoglycerate and phosphoenolpyruvate (PEP), was induced at 200 µmol photons m⁻² s⁻¹ in *LGS* while being reduced in the other strains. This pattern is consistent with diversion of carbon not committed to glycogen through PEP and acetyl-CoA toward lipid biosynthesis. Rre37, induced specifically in *LGS* and most strongly at 300 µmol photons m⁻² s⁻¹, regulates glycolytic and glycogen-catabolic genes (Azuma et al., 2011). As a 2-OG/NtcA-responsive regulator, its induction also links the *LGS* response to the carbon– nitrogen regulatory node affected by EFE expression. Together with the low level of extracellular overflow in *LGS* (Fig. 6), these data indicate that *LGS* preferentially accommodates surplus carbon through intracellular rerouting and storage rather than through secretion of organic acids.

## 4. Discussion

### 4.1 The 2-OG node determines whether increased photosynthetic flux is converted into product or redirected elsewhere

The growth, photosynthetic performance, and proteome allocation patterns across the tested strains point to a common metabolic determinant: the intracellular balance of 2-OG. In cyanobacteria, 2-OG integrates carbon and nitrogen metabolism by linking TCA cycle carbon flux, GS–GOGAT-mediated nitrogen assimilation and arginine biosynthesis, while also serving as a key metabolic signal sensed by the PII protein and the master nitrogen regulator NtcA (Llácer et al., 2010; Forchhammer and Selim, 2020). In the engineered strains, 2-OG is additionally a direct substrate of EFE (Eckert et al., 2014). The results of *ΔglgC+EFE* therefore show that increased availability of 2-OG does not automatically translate into proportionally higher ethylene production. Rather, continuous withdrawal of 2-OG by EFE perturbs the balance between carbon supply, nitrogen assimilation, and product formation. The effect becomes more pronounced with increasing irradiance and parallels the threshold-like growth collapse observed above a critical PFD (Fig. 1A). Thus, the capacity to maintain carbon–nitrogen and redox homeostasis, rather than precursor availability alone, appears to determine how effectively surplus photosynthetic flux can be converted into product.

### 4.2 Carbon-sink effects increase electron supply, but electron availability does not guarantee product formation

The failure of increased electron transport to translate proportionally into ethylene production in the EFE-containing strains is also consistent with their metabolic stoichiometry. Operation of the CBB cycle requires an ATP:NADPH ratio that linear electron transport alone cannot supply, making cyclic and other alternative electron flows necessary to balance cellular energy metabolism (Knoop and Steuer, 2015; Nikkanen et al., 2021). EFE couples 2-OG consumption to arginine hydroxylation and therefore provides a sink for both carbon and reducing equivalents (Fukuda et al., 1992). However, the stoichiometry of EFE does not match the full NADPH surplus associated with loss of glycogen synthesis (Table 1-2). Consequently, the additional electron flow cannot be expected to be channelled entirely into ethylene production. The remaining reductant must be accommodated by alternative electron sinks, overflow metabolism, or reduced overall carbon fixation. The present data support a contribution from overflow metabolism, whereas the precise contribution of alternative electron-flow pathways cannot be resolved from the proteomic dataset alone.

### 4.3 2-OG withdrawal reveals a phosphate constraint at the GS-GOGAT node

Introducing EFE into the *ΔglgC* background was associated with a secondary Pi limitation despite replete external phosphate. This is consistent with competition between EFE- mediated 2-OG consumption and GS-GOGAT-dependent nitrogen assimilation (Veetil et al., 2017): reduced GS-GOGAT flux would constrain glutamine supply for nucleotide and amino-acid synthesis, and the accompanying shift toward nucleotide turnover and cellular repair would increase reliance on internal phosphate recycling. The proteome of *ΔglgC+EFE* matches this expectation - induction of the Pst1 and Pst2 transport systems and of exopolyphosphatase Ppx, alongside near-WT polyphosphate kinase and reduced ATP synthase (Fig. 5B; Supplementary Table S1) indicate mobilisation of polyphosphate as an emergency Pi source (Veetil et al., 2017; Sanz-Luque et al., 2020).

An alternative possibility is that the same proteomic pattern reflects increased flux through the PPK/Ppx couple operating as a futile cycle, in which ATP is consumed during polyphosphate synthesis and Pi is released upon its degradation. However, the reduced abundance of ATP synthase in *ΔglgC+EFE* is not consistent with this expectation, and is instead consistent with limited availability of Pi, which is a substrate of ATP synthase. A contribution of polyphosphate cycling can nevertheless not be excluded on the basis of protein abundances alone. This distinction is relevant for strain design, as deletion of PPK increases photosynthetic productivity in *Synechocystis* (Sebesta et al., 2024), which is consistent both with removal of an ATP-consuming cycle and with altered Pi buffering. Direct quantification of intracellular Pi and polyphosphate pools would be required to determine whether Pi mobilisation or polyphosphate cycling predominates.

Ethylene itself is a signal in *Synechocystis*, sensed by SynEtr1 (Slr1212, also known as UirS), a hybrid light- and ethylene-sensing histidine kinase (Lacey and Binder, 2016), so induction of phosphate transporters might in principle reflect ethylene signalling rather than phosphate status. However, this appears unlikely here: the transporters upregulated by exogenous ethylene are those for chloride, manganese and amino acids rather than phosphate (Lacey et al., 2018), ethylene produced by EFE affects far fewer transcripts, mainly *csiR1* and *slr1214* (Kuchmina et al., 2017), and ethylene production rates were comparable in *EFE* and *ΔglgC+EFE* (Fig. 1B) whereas the phosphate response was specific to *ΔglgC+EFE*.

*ΔglgC+EFE* therefore appears to respond to the 2-OG drain not simply by reducing product formation, but by reallocating resources toward maintenance of phosphate homeostasis. This identifies the 2-OG/GS-GOGAT/Pi axis as an important potential constraint on heterologous pathways that withdraw TCA-cycle intermediates. From a strain-engineering perspective, productivity may therefore be improved either by reducing the rate of 2-OG withdrawal to a level compatible with GS-GOGAT capacity or by increasing the ability of the cell to maintain phosphate homeostasis under the imposed demand. The resulting picture of carbon, energy and phosphate allocation in *ΔglgC+EFE* under high irradiance is summarised in Fig. 7.

**Fig. 7.**
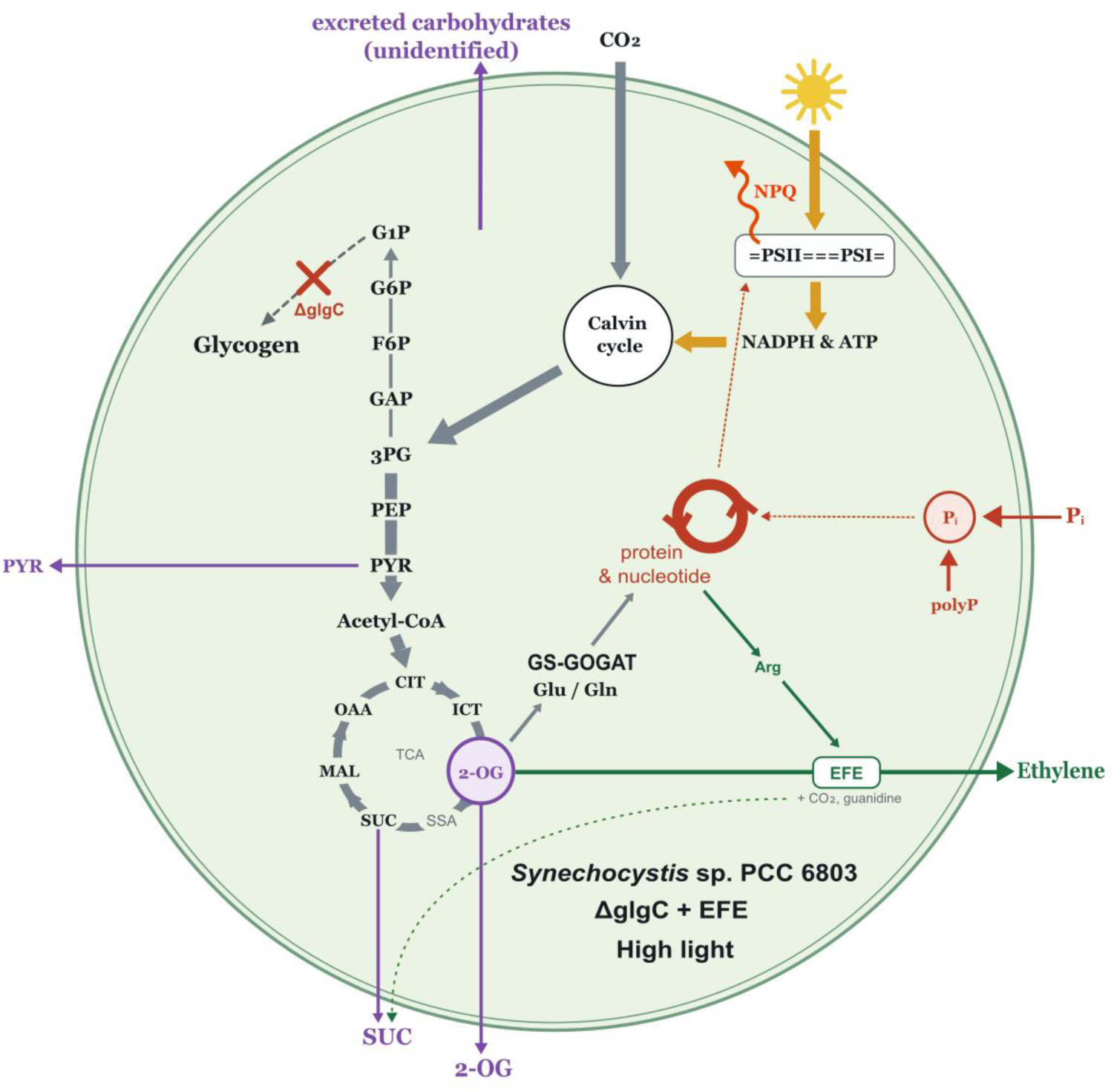
Proposed allocation of carbon, energy and phosphate in *Synechocystis* sp. PCC 6803 *ΔglgC+EFE* under high irradiance. Blocking glycogen synthesis (*ΔglgC*, red cross) prevents storage of fixed carbon as glycogen and redirects carbon from the Calvin–Benson–Bassham cycle through lower glycolysis and the TCA cycle (grey arrows). Light energy captured by the photosynthetic electron transport chain supplies NADPH and ATP (gold arrows), with excess excitation dissipated by non-photochemical quenching (NPQ, red). Heterologous EFE consumes 2-oxoglutarate (2-OG) together with arginine to produce ethylene, CO_2_, guanidine and succinate (green arrows). The resulting 2-OG withdrawal competes with the GS-GOGAT cycle for substrate, constraining glutamine and glutamate supply for protein and nucleotide synthesis and thereby increasing demand for inorganic phosphate (Pi). Increased Pi demand is accompanied by induction of the Pst transport systems and by mobilisation of polyphosphate (polyP) through exopolyphosphatase (red). Carbon not accommodated by these routes is excreted as pyruvate (PYR), 2-OG, succinate (SUC) and unidentified carbohydrates (purple arrows). Dotted arrows indicate proposed links inferred from proteomic and fluorescence data rather than measured fluxes; intracellular Pi and polyP pools were not quantified in this study. Abbreviations: 3PG, 3-phosphoglycerate; Acetyl-CoA, acetyl coenzyme A; Arg, arginine; CIT, citrate; F6P, fructose-6-phosphate; G1P, glucose-1-phosphate; G6P, glucose-6- phosphate; GAP, glyceraldehyde-3-phosphate; Gln, glutamine; Glu, glutamate; ICT, isocitrate; MAL, malate; OAA, oxaloacetate; PEP, phosphoenolpyruvate; PSI/PSII, photosystem I/II; SSA, succinic semialdehyde.

### 4.4 Photoprotection provides a readout of cellular energy economy

The tested strains also adopted distinct photoprotective strategies, which reflected different metabolic states. *ΔglgC* preferentially reduced PSII excitation through a pronounced State 1→2 transition while maintaining low OCP-mediated NPQ, whereas *ΔglgC+EFE* showed strong OCP induction, PBS reduction, and a State 2→1 transition (Figures 2 and 5). State transitions redirect rather than dissipate excitation energy and are energetically inexpensive, representing a preferred response when biosynthetic capacity is constrained (Calzadilla and Kirilovsky, 2020). By contrast, OCP-mediated NPQ dissipates absorbed excitation as heat and requires synthesis and turnover of the photoprotective machinery. The strong NPQ response of *ΔglgC+EFE* at high irradiance is therefore consistent with a strain experiencing greater excitation pressure while simultaneously facing a metabolic constraint on biosynthetic capacity.

LGS employs a third route, selectively reducing PBS subunits together with induction of the rod-linker protein CpcC2, without engaging either state transitions or OCP-mediated NPQ to the same extent as *ΔglgC+EFE*. Coexistence of these distinct responses suggests that cyanobacteria can redistribute excitation through several partially substitutable mechanisms, with the relative contribution of each pathway depending on the prevailing metabolic state. Photoprotection may therefore serve not only as a response to excess light but also as an indirect readout of the cellular capacity to consume the resulting energy and reductant.

### 4.5 The fate of surplus electrons remains unresolved

Two mechanistic questions remain open. First, both NDH-1 and NDH-2 subunits were downregulated in *ΔglgC* (Supplementary Table S1), indicating altered capacity for cyclic electron flow. At the same time, flavodiiron proteins Flv1/Flv3, which catalyse a major alternative electron-flow pathway involving Mehler-like O_2_ reduction in cyanobacteria (Nikkanen et al., 2021) were not differentially regulated. The current proteomic data therefore do not identify a specific pathway responsible for disposal of the surplus reductant. Possible explanations include post-translational activation of flavodiiron proteins or terminal oxidases

operating below saturation (Lea-Smith et al., 2013), as well as overflow excretion acting as a major downstream sink.

Second, strong induction of carboxysome shell proteins CcmL and CcmN in *ΔglgC+EFE* (Supplementary Table S1) raises the possibility that phosphate stress also affects the CO_2_- concentrating mechanism, although bicarbonate-uptake systems showed no consistent changes. Quantification of dissolved O_2_ and CO_2_ exchange, for example by membrane-inlet mass spectrometry (Müller et al., 2019), would distinguish between alternative mechanisms of reductant disposal and determine whether impaired inorganic-carbon acquisition contributes to the phenotype.

### 4.6 Glycogen-overflow phenotypes are strain-specific

Comparison with other cyanobacteria strains further indicates that the consequences of glycogen deletion depend on strain-specific regulatory architecture rather than following a universal pattern. *Synechocystis* sp. PCC 6803 *ΔglgC* predominantly excretes 2-OG and pyruvate (Carrieri et al., 2015; Cano et al., 2018), whereas *Synechococcus elongatus* PCC 7942 *ΔglgC* releases glutamate as the major overflow product under nitrogen-replete conditions (Kato et al., 2024). *Synechococcus* sp. PCC 7002 *ΔglgC* produces a broader mixture of organic acids under nitrogen deprivation (Davies et al., 2014; Jackson et al., 2015). These different overflow signatures imply that each species partitions excess carbon among 2-OG excretion, glutamate accumulation, GS-GOGAT flux, and other sinks according to its own regulatory architecture. The Pi-limitation phenotype identified here may therefore not be reproduced quantitatively in other cyanobacterial chassis. More broadly, the comparison supports the view that the consequences of deleting a storage pathway depend on the surrounding network and regulatory context rather than on the deleted reaction alone (Xiong et al., 2017).

An additional, unidentified carbohydrate-like compound was consistently excreted by *ΔglgC* and *ΔglgC+EFE*, but not by *EFE* alone, pointing to a glycogen-loss-driven and EFE- independent overflow route. To our knowledge, this phenotype has not been reported previously: previous studies of *ΔglgC* mutants have described overflow of organic acids and amino acids, including 2-OG, pyruvate, succinate, malate, and glutamate, but not carbohydrate-like compounds (Cano et al., 2018; Cantrell et al., 2023; Kato et al., 2024; Alford et al., 2025). Its chemical identity remains unresolved; possible sources include leakage of the CBB cycle sugar phosphates or products of polysaccharide turnover. Together with the low extracellular overflow observed in *LGS*, this finding suggests that loss of glycogen synthesis distributes surplus carbon across several partially overlapping routes rather than a single dominant overflow pathway.

## 5. Conclusions

Taken together, the present results identify the 2-OG/GS-GOGAT/Pi axis as a central constraint on the conversion of surplus photosynthetic flux into heterologous products (Fig. 7). The key limitation is not simply the availability of carbon or reducing equivalents, but the ability of the cell to maintain carbon–nitrogen, phosphate, and energetic homeostasis while sustaining product flux. This is particularly evident in *ΔglgC+EFE*, where continuous 2-OG withdrawal is associated with a phosphate-stress response, enhanced photoprotection, and reduced growth despite increased photosynthetic electron transport. The results further show that removal of a carbon-storage pathway does not produce a single, predictable redistribution of flux. The metabolic consequences of carbon-sink removal emerge from the interaction between the engineered pathway and the broader regulatory network. Productive cyanobacterial chassis should be therefore designed not only around the catalytic capacity of the target pathway, but also around the cellular processes that sustain its substrate and cofactor supply. For pathways that withdraw 2-OG or related TCA-cycle intermediates, phosphate homeostasis and the capacity of the GS-GOGAT network may constitute important design constraints that should be considered at the engineering stage.

## Data availability statement

The mass spectrometry proteomics data generated in this study are available via the ProteomeXchange Consortium and the PRIDE partner repository under the dataset identifier PXD076169. The chlorophyll-fluorescence analysis pipeline, available at https://www.cyano.tools/, is archived at https://doi.org/10.5281/zenodo.21916564. All other data are included in the article and its Supplementary Material.

## Author Contributions

TZa: Conceptualization, Investigation, Methodology, Validation, Software, Data curation, Formal Analysis, Visualization, Supervision, Project administration, Funding acquisition, Resources, Writing – original draft, Writing – review & editing. JS: Investigation, Methodology, Validation, Resources, Writing – review & editing. HL: Investigation, Methodology, Validation, Writing – review & editing. TZl: Methodology, Validation, Data curation, Writing – review & editing. TD: Investigation, Methodology, Writing – review & editing. RSa: Investigation, Methodology, Validation, Software, Formal Analysis, Data curation, Visualization, Writing – review & editing. DB: Methodology, Writing – review & editing. MČ, MB, ML: Investigation, Methodology, Data curation, Formal Analysis, Writing – review & editing. RH and RSt: Investigation, Methodology, Validation, Software, Formal Analysis, Writing – review & editing. JČ and JY: Conceptualization, Investigation, Funding acquisition, Writing – review & editing.

## Generative AI statement

The authors declare that generative AI was used in the creation of this manuscript. Anthropic’s Claude was used to assist with data analysis and statistical evaluation and with drafting and editing of the manuscript text. All AI-assisted analyses and text were reviewed, verified against the primary data, and edited by the authors, who take full responsibility for the content of the published article.

