## Supplementary material for "Glycogen removal enhances ethylene production in *Synechocystis* sp. PCC 6803 but exposes a 2-oxoglutarate–GS-GOGAT–phosphate constraint"

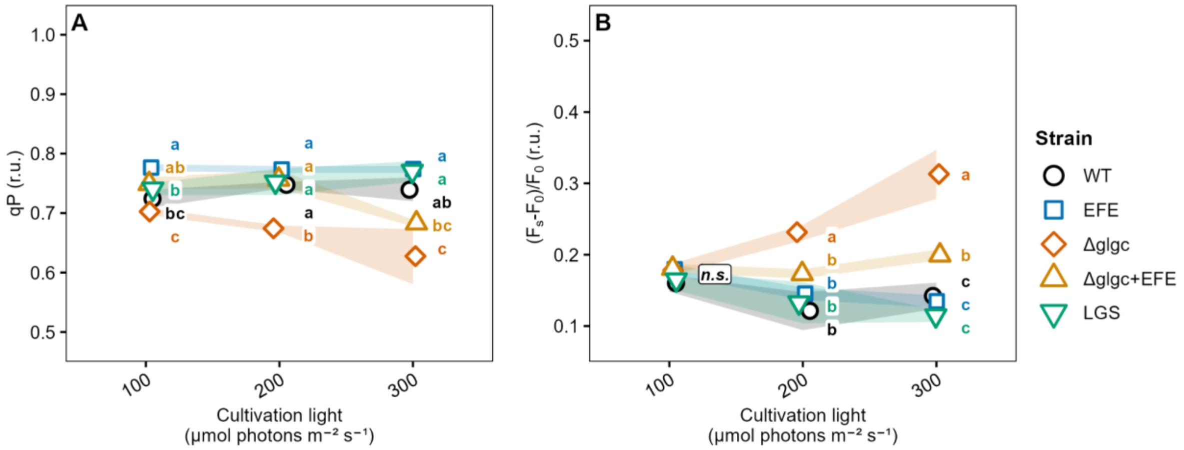


**Figure S1** Additional chlorophyll-fluorescence parameters derived from AquaPen NPQ3 measurements in the tested *Synechocystis* strains cultivated at 100–300 µmol photons m$^{-2}$ s$^{-1}$, complementing Fig. 2. **(A)** Coefficient of photochemical quenching (qP; Eq. 3), reflecting the fraction of open PSII reaction centres. **(B)** Relative rise of steady-state fluorescence above the dark-acclimated minimum, (Fs-FO)/FO, used as an indicator of PSII excitation pressure under actinic illumination. Both parameters were determined after full light acclimation, designated “Lss” by the AquaPen software. Symbols show mean values and ribbons the corresponding standard deviations. Letters indicate significant differences between strains within each light intensity (one-way ANOVA followed by Tukey’s HSD, $p < 0.05$); n.s., no significant pairwise differences at the indicated light intensity; n = 3.

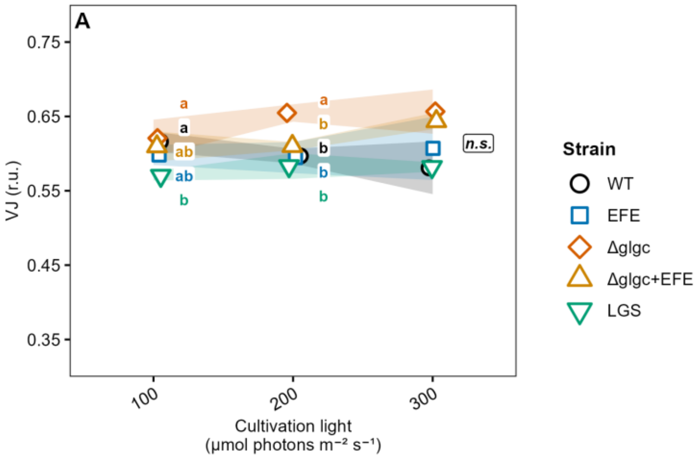


**Figure S2** Relative variable fluorescence at the J step ($V_J$), derived from the fast fluorescence-induction (OJIP) transients of the tested *Synechocystis* strains cultivated at 100–300 µmol photons $m^{-2}$ $s^{-1}$, complementing Fig. 4. $V_J$ was calculated according to Eq. 8 from the fluorescence level at 2 ms and reports the redox state of the plastoquinone pool, with higher values indicating a more reduced pool. Measurements were performed after 15 min of dark acclimation. Symbols show mean values and ribbons the corresponding standard deviations. Letters indicate significant differences between strains within each light intensity (one-way ANOVA followed by Tukey's HSD, $p < 0.05$); n.s., no significant pairwise differences at the indicated light intensity; $n = 3$.

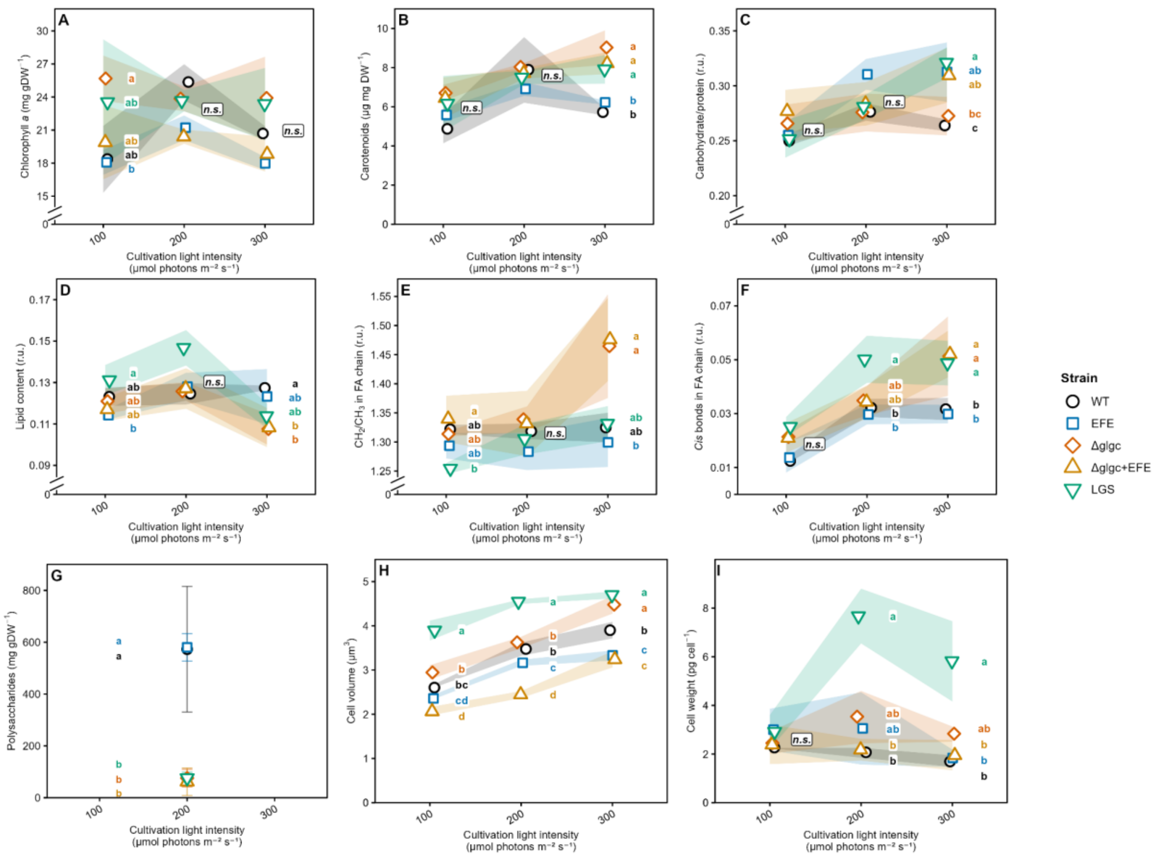


**Figure S3** Pigment content, macromolecular composition and cell parameters of the tested *Synechocystis* strains cultivated at 100–300 µmol photons $m^{-2}$ $s^{-1}$. **(A)** Chlorophyll *a* and **(B)** total carotenoid content per unit dry weight, determined spectrophotometrically. **(C–F)** Macromolecular composition determined by FTIR spectroscopy: **(C)** carbohydrate-to-protein ratio, from the band amplitudes at ~1152 $cm^{-1}$ and ~1652 $cm^{-1}$; **(D)** lipid content, from the band at ~1735 $cm^{-1}$ normalised to the protein band at ~1652 $cm^{-1}$; **(E)** ratio of the asymmetric $CH_2$ and $CH_3$ stretching amplitudes (~2925 and ~2960 $cm^{-1}$), used as a proxy for the relative fatty-acid chain length, with higher values corresponding to longer chains; **(F)** relative content of *cis* double bonds in the fatty-acid chains. **(G)** Polysaccharide content, determined only for cultures grown at 200 µmol photons $m^{-2}$ $s^{-1}$; the method does not fully distinguish glycogen from other cellular polysaccharides. **(H)** Mean cell volume and **(I)** dry weight per cell. Symbols show averages; ribbons (A–F, H, I) and error bars (G) show the corresponding standard deviations. Letters indicate significant differences between strains within each light intensity ($p < 0.05$); n.s., no significant pairwise differences at the indicated light intensity; n = 3-4.

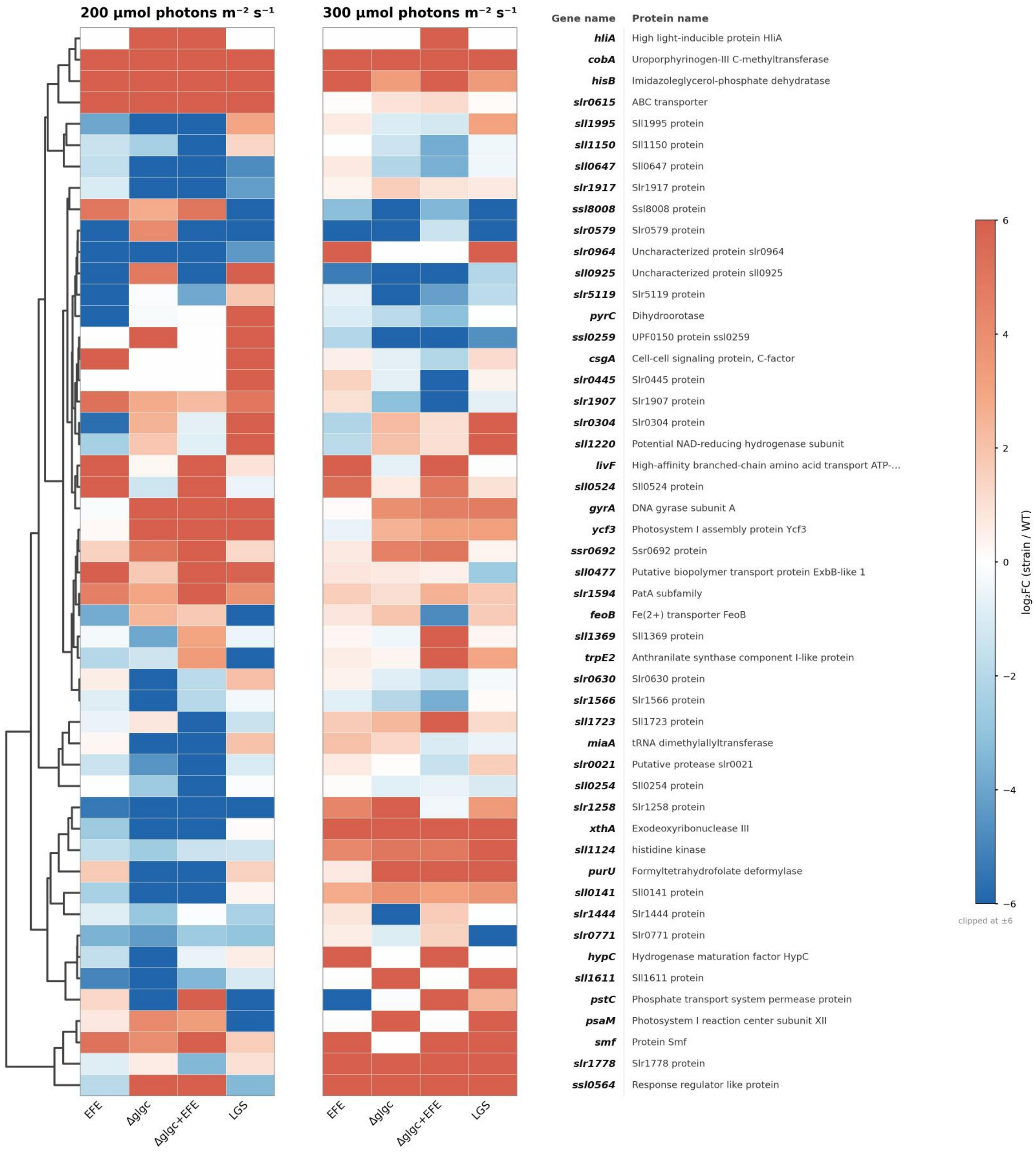


**Figure S4** Top 50 proteins showing the largest abundance change between WT and the tested *Synechocystis* strains at 200 and 300 µmol photons m$^{-2}$ s$^{-1}$. Colour encodes the $\log_2$ fold change of each strain against WT, calculated per biological replicate as $\log_2$(replicate intensity / mean WT intensity) and averaged over four biological replicates, with the two light conditions normalised independently. Rows were ordered by hierarchical clustering (dendrogram on the left) and are annotated with the gene name and the corresponding protein name. White cells indicate proteins that were either identical to WT, or were not quantified in the given strain and light condition. Only proteins with at least two unique peptides, or one unique peptide and at least 10 peptides measured, were considered. The complete dataset and the assignment of proteins to functional categories are provided in Supplementary Table S1.

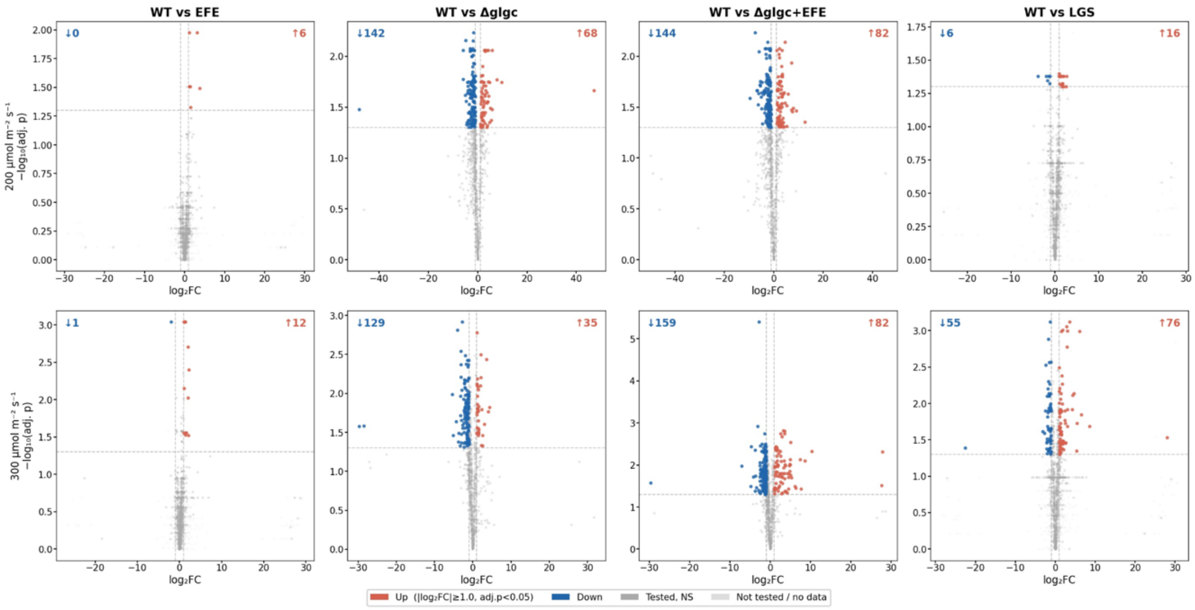


**Figure S5** Volcano plots of the complete proteomic dataset for each tested *Synechocystis* strain relative to WT at 200 (upper row) and 300 µmol photons $m^{-2}$ $s^{-1}$ (lower row). The x-axis shows the $\log_2$ fold change of the given strain against WT and the y-axis the $-\log_{10}$ of the Benjamini–Hochberg-adjusted p-value. Dashed lines mark the thresholds used throughout the study ($|\log_2 FC| \geq 1.0$ and adjusted $p < 0.05$). Proteins passing both thresholds are shown in red (increased abundance relative to WT) or blue (decreased abundance), and their numbers are given in the upper corners of each panel; dark grey denotes proteins that were tested but not significantly changed, and light grey proteins for which no test could be performed because they were quantified in only one of the two compared groups — these appear at extreme $\log_2$ fold changes. Four biological replicates per strain; the two light conditions were normalised independently. Note that the axis ranges differ between panels.

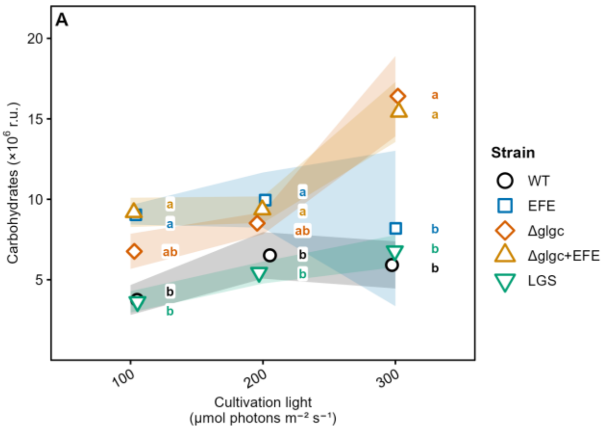


**Figure S6** Unidentified overflow carbohydrate measured in the cultivation medium of the tested *Synechocystis* strains at 100-300 µmol photons $m^{-2}$ $s^{-1}$, complementing Fig. 6. Values are expressed relative to ribitol as internal standard. Symbols show mean values and ribbons the corresponding standard deviations; n = 3-4. Letters indicate significant differences between strains within each light intensity ($p < 0.05$).

**Table S1** Complete quantitative proteomic dataset for the tested *Synechocystis* strains cultivated at 200 and 300 µmol photons $m^{-2}$ $s^{-1}$, together with the assignment of proteins to the 25 predefined functional categories used in Fig. 5B. Protein abundances were determined by label-free quantification; 438 of the 2 368 detected proteins were assigned to a functional category. The categories were selected a priori for their established relevance to the strains studied and cover photosynthesis, carbon fixation, central carbon metabolism, nitrogen assimilation, carbon storage sinks, inorganic nutrient and metal-ion transport, amino-acid transport and biosynthesis, two-component signalling, chlorophyll biosynthesis, DNA repair, and purine and pyrimidine metabolism. $Log_2$ fold changes were computed for each biological replicate as $log_2$(replicate intensity / mean WT intensity), with the 200 and 300 µmol photons $m^{-2}$ $s^{-1}$ conditions normalised independently; proteins with a missing value in a given replicate were excluded from the corresponding median rather than imputed. Category-level values were summarised per replicate as the median $log_2$ fold change across all proteins assigned to that category, yielding four biological replicates per strain and category. WT-noise values were obtained by applying the identical calculation to each individual WT replicate against the mean WT intensity, so that any deviation from zero reflects within-WT replicate variability and provides a baseline against which the fold changes of the other strains can be assessed. Only proteins with at least two unique peptides, or one unique peptide and at least 10 peptides measured, were retained. The mass spectrometry proteomics data are available via the ProteomeXchange Consortium and the PRIDE partner repository under the dataset identifier PXD076169.

| Gene | Accession | Protein name | Log2FC at 200 µE m⁻² s⁻¹ (rel. to WT) EFE | Δglgc | Δglgc+EFE | LGS | Log2FC at 300 µE m⁻² s⁻¹ (rel. to WT) EFE | Δglgc | Δglgc+EFE | LGS |
|---|---|---|---|---|---|---|---|---|---|---|
| **Phycobilisomes (PBS) (n=15)** | | | | | | | | | | |
| *cpcC2* | P73204 | Phycobilisome 32.1 kDa linker polypeptide, phycocyanin-associated, rod 2 | -0.17 | -0.56 | -0.60 | -0.19 | 0.02 | -0.37 | -0.80 | 4.21 |
| *cpcF* | P72652 | Phycocyanin alpha phycocyanobilin lyase CpcF | 0.39 | -1.76 | -1.02 | 0.63 | 0.49 | -0.33 | -1.43 | -0.25 |
| *apcB* | Q01952 | Allophycocyanin beta chain | -0.71 | 0.31 | -0.17 | 0.34 | -0.45 | 0.02 | -0.48 | -1.14 |
| *apcE* | Q55544 | Phycobiliprotein ApcE | -0.33 | -0.36 | -0.72 | 0.20 | -0.13 | -0.08 | -0.93 | -0.24 |
| *cpcG* | P73093 | Phycobilisome rod-core linker polypeptide CpcG | -0.25 | -0.37 | -0.70 | 0.22 | -0.18 | -0.08 | -0.88 | -0.17 |
| *cpcE* | P73638 | Phycocyanobilin lyase subunit alpha | 0.30 | -0.34 | 0.21 | 0.87 | 0.16 | -0.38 | -0.15 | -0.19 |
| *cpcA* | Q54715 | C-phycocyanin alpha subunit | -0.86 | -0.11 | -0.66 | 0.13 | -0.62 | 0.08 | -0.63 | -0.82 |
| *cpcC1* | P73203 | Phycobilisome 32.1 kDa linker polypeptide, phycocyanin-associated, rod 1 | -0.43 | -0.83 | -0.80 | -0.30 | 0.05 | 0.08 | -0.34 | -0.26 |
| *apcA* | Q01951 | Allophycocyanin alpha chain | -0.26 | 0.55 | -0.10 | 0.02 | -0.17 | 0.74 | -0.14 | 0.02 |
| *cpcB* | Q54714 | C-phycocyanin beta subunit | -0.30 | -0.13 | -0.64 | -0.15 | -0.13 | 0.43 | -0.52 | -0.17 |
| *cpcD* | P73202 | Phycobilisome 8.9 kDa linker polypeptide, phycocyanin-associated, rod | 0.03 | 0.58 | 0.13 | -0.30 | -0.48 | 0.63 | 0.11 | -0.13 |
| *apcF* | P74551 | Allophycocyanin subunit beta-18 | -0.13 | 0.50 | 0.45 | 0.34 | -0.20 | 0.62 | 0.34 | -0.48 |
| *apcD* | P72870 | Allophycocyanin subunit alpha-B | -0.16 | -0.17 | -0.42 | 0.35 | -0.07 | 0.15 | -0.53 | -0.44 |
| *apcC* | Q01950 | Phycobilisome 7.8 kDa linker polypeptide, allophycocyanin-associated, core | 0.45 | -0.24 | -0.01 | 0.24 | 0.53 | -0.10 | 0.07 | 0.13 |
| *pcyA* | Q55891 | Phycocyanobilin:ferredoxin oxidoreductase | 0.06 | -0.34 | -0.46 | 0.40 | 0.20 | 0.31 | -0.47 | -0.07 |
| **Photosynthetic electron transport (PSET) (n=58)** | | | | | | | | | | |
| *psbE* | P09190 | Cytochrome b559 subunit alpha | -8.40 | -7.58 | -7.79 | -8.80 | -0.34 | -0.81 | -0.28 | -0.80 |
| *petF* | P27320 | Ferredoxin-1 | -0.15 | -0.81 | 0.98 | 0.12 | 3.15 | 4.92 | 5.21 | 5.95 |
| *atpF* | P27181 | ATP synthase subunit b | 0.23 | 1.17 | 5.45 | 0.27 | -0.27 | 0.76 | 0.21 | 0.24 |
| *atpA* | P27179 | ATP synthase subunit alpha | -0.38 | -0.24 | -0.64 | -0.13 | -4.63 | -4.73 | -5.34 | -4.36 |
| *psbV* | Q55013 | Photosystem II extrinsic protein V | 0.13 | 0.62 | -0.16 | 0.11 | -0.51 | 0.99 | 5.27 | -0.11 |
| *psaK1* | P72712 | Photosystem I reaction center subunit PsaK 1 | -4.67 | -4.73 | -3.28 | -1.04 | -1.26 | -0.70 | -0.51 | -0.83 |
| *ycf3* | P74063 | Photosystem I assembly protein Ycf3 | 0.12 | 4.55 | 4.20 | 3.94 | -0.56 | 2.54 | 3.13 | 3.17 |
| *psbL* | Q55354 | Photosystem II reaction center protein L | 2.22 | 2.76 | 3.42 | 2.01 | -0.42 | -0.97 | 0.04 | -1.63 |
| *psbY* | P73676 | Photosystem II reaction center protein Y | | | | | -0.47 | 3.33 | -0.56 | -2.25 |
| *btpA* | P72966 | Photosystem I biogenesis protein BtpA | -1.14 | -1.76 | -2.99 | -0.85 | 1.25 | 0.51 | -0.72 | 1.31 |
| *atpE* | P27182 | ATP synthase subunit c | -1.42 | 1.22 | 1.89 | 0.83 | -1.04 | 2.19 | 1.93 | 2.88 |
| *atpB* | P27178 | ATP synthase subunit a | -0.71 | -2.26 | -2.62 | 0.55 | -0.08 | -0.17 | -0.89 | -0.47 |
| *ssl2559* | P73171 | Ferredoxin | -1.01 | -2.29 | -1.20 | -1.19 | -0.16 | 0.77 | 0.53 | 0.89 |
| *slr2059* | P73811 | Ferredoxin | -0.58 | -0.62 | -1.33 | -0.85 | 1.59 | 2.27 | 1.18 | 1.40 |
| *ycf48* | P73069 | Photosystem II assembly lipoprotein Ycf48 | 0.29 | 0.35 | -0.89 | -0.02 | 1.71 | 0.48 | 2.06 | 1.79 |
| *petJ* | P46445 | Cytochrome c6 | 0.09 | 2.01 | 0.92 | 1.27 | -0.71 | -0.03 | -0.70 | -1.04 |
| *psbH* | P14835 | Photosystem II reaction center protein H | -0.06 | 1.89 | 1.34 | 1.16 | -0.44 | 0.67 | 1.02 | -1.24 |
| *sll0662* | Q55980 | Ferredoxin | 0.85 | 1.15 | 1.25 | 1.44 | 1.63 | 1.85 | -0.78 | 1.05 |
| *psbF* | P09191 | Cytochrome b559 subunit beta | 0.15 | 1.52 | 1.32 | 0.89 | -0.58 | -0.07 | 0.64 | -0.76 |
| *psaK2* | P74564 | Photosystem I reaction center subunit PsaK 2 | 0.82 | 1.43 | 1.51 | 1.45 | 0.09 | 0.29 | 0.71 | 0.43 |
| *petB* | Q57038 | Cytochrome b6 | -0.30 | -0.03 | 0.00 | 0.98 | -0.12 | -0.36 | -1.45 | 0.16 |
| *petD* | P27589 | Cytochrome b6-f complex subunit 4 | -0.36 | 0.55 | 0.41 | 1.43 | -0.61 | -0.92 | -0.38 | -0.46 |
| *psaM* | P72986 | Photosystem I reaction center subunit XII | 0.43 | 1.43 | 1.37 | | | | | |
| *psaC* | P32422 | Photosystem I iron-sulfur center | -0.12 | 0.57 | 0.09 | 0.12 | -0.02 | 1.41 | 0.49 | 0.56 |
| *slr1739* | P73731 | Photosystem II reaction center Psb28 protein | 0.89 | 1.37 | 1.31 | 0.51 | 0.19 | 0.77 | 0.79 | 0.11 |
| *psb34* | P74067 | Photosystem II assembly protein Psb34 | 0.15 | 0.16 | 0.19 | -0.85 | -1.05 | -1.26 | -0.68 | -1.31 |
| *psb27* | P74367 | Photosystem II lipoprotein Psb27 | 0.47 | 0.90 | 0.58 | 0.46 | 0.35 | 1.27 | 0.67 | 0.53 |
| *psaA* | P29254 | Photosystem I P700 chlorophyll a apoprotein A1 | -0.61 | 0.88 | -0.07 | 0.88 | -0.28 | -0.37 | -0.59 | -1.22 |
| *atpG* | P17253 | ATP synthase gamma chain | -0.10 | -0.62 | -0.68 | 0.28 | -0.08 | -0.35 | -1.19 | 0.38 |
| *petF* | P74449 | Ferredoxin | -0.33 | 0.61 | 0.03 | 1.09 | -0.50 | 0.80 | 0.42 | 0.05 |
| *cpcL* | P74625 | Photosystem I-associated linker protein CpcL | -0.05 | -0.27 | -0.21 | -0.12 | -0.25 | -0.70 | -1.08 | -0.63 |
| *atpH* | P27180 | ATP synthase subunit delta | -0.11 | -0.72 | -0.43 | 0.63 | 0.07 | -0.55 | -1.07 | -0.01 |
| *psbU* | Q55332 | Photosystem II extrinsic protein U | -0.25 | 0.39 | -0.08 | 0.02 | -0.53 | -0.88 | -0.74 | -1.01 |
| *petP* | P72798 | Cytochrome b6-f regulatory subunit PetP | -0.10 | 0.99 | 0.08 | 0.49 | -0.32 | 0.75 | 0.40 | 0.51 |
| *petE* | P21697 | Plastocyanin | 0.23 | 0.00 | 0.08 | -0.42 | 0.02 | 0.92 | 0.63 | 0.65 |
| *petF* | P73388 | Ferredoxin | 0.76 | 0.31 | 0.28 | -0.17 | -0.10 | -0.90 | -0.29 | -0.52 |
| *psaB* | P29255 | Photosystem I P700 chlorophyll a apoprotein A2 | 0.12 | -0.37 | -0.51 | 0.66 | -0.04 | -0.14 | -0.84 | -0.30 |
| *atpC* | P26533 | ATP synthase epsilon chain | -0.20 | -0.63 | -0.75 | 0.11 | 0.42 | 0.73 | -0.21 | 0.84 |
| *atpF2* | P27183 | ATP synthase subunit b' | 0.39 | 0.83 | 0.46 | 0.24 | -0.31 | -0.06 | -0.27 | -0.24 |
| *slr1205* | P73355 | Ferredoxin component | 0.63 | 0.12 | 0.71 | 0.83 | -0.05 | 0.18 | 0.41 | -0.07 |
| *psaD* | P19569 | Photosystem I reaction center subunit II | 0.03 | -0.57 | -0.58 | 0.47 | -0.27 | -0.21 | -0.81 | -0.21 |
| *psaE* | P12975 | Photosystem I reaction center subunit IV | -0.23 | -0.46 | -0.80 | -0.05 | -0.22 | 0.48 | -0.05 | 0.23 |
| *atpD* | P26527 | ATP synthase subunit beta | 0.01 | 0.00 | -0.78 | 0.18 | -0.24 | 0.45 | -0.24 | 0.03 |
| *ycf4* | P72705 | Photosystem I assembly protein Ycf4 | 0.09 | -0.01 | -0.49 | 0.77 | 0.70 | 0.30 | -0.65 | 0.74 |
| *psaL* | P37277 | Photosystem I reaction center subunit XI | 0.01 | -0.31 | -0.43 | 0.04 | 0.01 | 0.77 | 0.36 | 0.29 |
| *petH* | Q55318 | Ferredoxin–NADP reductase | 0.01 | -0.43 | -0.51 | 0.53 | -0.18 | -0.06 | -0.76 | 0.12 |
| *petC1* | P26290 | Cytochrome b6-f complex iron-sulfur subunit 1 | 0.11 | 0.76 | 0.46 | 0.61 | -0.13 | 0.66 | -0.02 | 0.35 |
| *petF* | P74159 | Ferredoxin | 0.69 | 0.46 | 0.43 | 0.25 | 0.18 | 0.64 | 0.63 | 0.30 |
| *psbD* | P09192 | Photosystem II D2 protein | -0.09 | -0.68 | -0.68 | 0.33 | -0.10 | -0.05 | -0.51 | -0.28 |
| *psbA2* | P16033 | Photosystem II protein D1 2 | -0.04 | -0.39 | -0.34 | 0.20 | -0.04 | -0.23 | -0.67 | -0.31 |
| *slr0148* | P74447 | Ferredoxin | -0.25 | 0.25 | -0.48 | 0.15 | -0.52 | 0.67 | 0.13 | 0.47 |
| *psbB* | P05429 | Photosystem II CP47 reaction center protein | -0.47 | -0.29 | -0.50 | 0.18 | -0.30 | -0.22 | -0.51 | -0.57 |
| *petA* | P26287 | Cytochrome f | 0.05 | 0.31 | 0.11 | 0.53 | 0.16 | 0.34 | -0.54 | 0.41 |
| *psaF* | P29256 | Photosystem I reaction center subunit III | 0.00 | -0.08 | -0.13 | 0.42 | -0.13 | 0.10 | -0.45 | -0.12 |
| *ycf39* | P74429 | Photosystem II assembly factor Ycf39 | 0.08 | -0.10 | 0.01 | 0.27 | 0.44 | 0.24 | 0.03 | 0.17 |
| *psb28* | Q55356 | Photosystem II reaction center Psb28 protein | -0.14 | 0.32 | -0.33 | -0.17 | -0.36 | 0.38 | -0.10 | -0.04 |
| *psbO* | P10549 | Photosystem II extrinsic protein O | 0.01 | 0.03 | -0.35 | 0.22 | -0.07 | 0.28 | -0.21 | -0.07 |
| *psbC* | P09193 | Photosystem II CP43 reaction center protein | 0.03 | 0.13 | 0.01 | 0.04 | 0.09 | 0.07 | -0.33 | 0.29 |
| **Respiratory electron transport (RET) (n=30)** | | | | | | | | | | |
| *ctaD* | P73907 | cytochrome-c oxidase | -3.18 | -0.70 | 1.05 | -0.60 | 0.68 | -1.01 | 3.65 | 1.71 |
| *cydA* | P73159 | Cytochrome oxidase d subunit I | -2.56 | -2.22 | -2.17 | -1.56 | 0.47 | 0.03 | -0.58 | -0.19 |
| *ndh* | P74614 | NADH dehydrogenase | 0.45 | -0.94 | -0.89 | 0.11 | 0.14 | -1.39 | -2.39 | -0.36 |
| *sdhB* | P73723 | Fumarate reductase iron-sulfur subunit | -1.66 | -1.66 | -2.20 | -1.62 | -0.23 | 1.07 | -1.66 | -1.26 |
| *ndhH* | P27724 | NAD(P)H-quinone oxidoreductase subunit H | -0.59 | -1.31 | -0.92 | 0.02 | 0.25 | 2.09 | -0.18 | -0.64 |
| *ndhB* | P72714 | NAD(P)H-quinone oxidoreductase subunit 2 | 1.26 | 1.32 | 1.99 | 1.19 | 0.50 | -0.30 | 1.77 | -0.21 |
| *ndhO* | P74771 | NAD(P)H-quinone oxidoreductase subunit O | 1.31 | 0.15 | 1.95 | 0.24 | 0.91 | -1.41 | 0.33 | 0.11 |
| *ndhF* | Q55465 | NADH dehydrogenase subunit 5 | -0.83 | 1.81 | 0.24 | 1.52 | -0.88 | -0.88 | -0.07 | -0.41 |
| *ndhD2* | P72823 | NAD(P)H-quinone oxidoreductase chain 4-2 | 0.31 | 0.90 | 0.46 | 1.05 | 0.64 | 0.43 | -0.32 | 1.72 |
| *ndhJ* | P19125 | NAD(P)H-quinone oxidoreductase subunit J | 0.09 | -1.21 | 1.25 | -1.72 | -0.37 | 0.73 | 0.28 | -0.64 |
| *ndh* | P73739 | NADH:ubiquinone reductase (non-electrogenic) | 0.14 | -0.30 | -1.68 | 0.59 | 0.41 | -0.55 | -0.26 | 0.09 |
| *ndhD* | Q55464 | NADH dehydrogenase subunit 4 | 0.85 | -0.52 | 1.64 | 0.53 | -0.09 | 0.03 | -0.71 | -0.42 |
| *ndhG* | P26523 | NAD(P)H-quinone oxidoreductase chain 6 | -0.72 | -0.10 | -0.95 | -0.97 | 0.10 | 0.61 | -1.39 | 0.54 |
| *ndhA* | P26522 | NAD(P)H-quinone oxidoreductase subunit 1 | 0.25 | -1.10 | -0.52 | 0.55 | 0.09 | 0.16 | -1.28 | -1.39 |
| *ctaD* | Q06473 | Cytochrome c oxidase subunit 1 | 0.00 | -0.63 | -0.99 | 1.14 | -0.07 | -0.74 | -0.53 | -0.83 |
| *ndhF* | Q55429 | NAD(P)H-quinone oxidoreductase chain 5 | 0.34 | -0.29 | -0.23 | -0.18 | 0.24 | -1.13 | 0.23 | 0.06 |
| *ctaC* | Q06474 | Cytochrome c oxidase subunit 2 | 0.56 | 1.07 | 0.60 | 0.88 | 0.12 | 0.97 | 1.08 | -0.09 |
| *ndhD1* | P32421 | NAD(P)H-quinone oxidoreductase chain 4-1 | 0.45 | 0.76 | -0.20 | 0.85 | 0.27 | -0.45 | -0.88 | -0.12 |
| *ndhM* | P74338 | NAD(P)H-quinone oxidoreductase subunit M | -0.24 | 0.05 | -0.39 | -0.79 | -0.14 | 0.45 | 0.04 | 0.17 |
| *ndhI* | P26525 | NAD(P)H-quinone oxidoreductase subunit I | 0.41 | -0.61 | -0.34 | 0.47 | 0.13 | -0.28 | -0.76 | -0.11 |
| *frdA* | P73479 | Succinate dehydrogenase flavoprotein subunit | 0.30 | -0.72 | -0.22 | 0.05 | 0.59 | 0.21 | -0.27 | 0.48 |
| *ndhN* | P74069 | NAD(P)H-quinone oxidoreductase subunit N | -0.22 | -0.21 | 0.35 | -0.06 | 0.32 | -0.52 | -0.65 | 0.06 |
| *ctaC* | P74044 | Cytochrome c oxidase subunit 2 | -0.31 | -0.12 | 0.62 | 0.51 | 0.30 | 0.04 | 0.24 | 0.12 |
| *ndhK1* | P19050 | NAD(P)H-quinone oxidoreductase subunit K 1 | -0.12 | -0.41 | -0.24 | -0.04 | 0.52 | 0.08 | -0.62 | 0.50 |
| *ndbB* | P73735 | Demethylphylloquinone reductase NdbB (NDH-2) | 1.95 | 1.24 | 2.65 | 0.61 | 0.81 | -1.30 | 1.13 | -1.26 |
| *petA* | P26287 | Cytochrome f (PetA) | 0.05 | 0.31 | 0.11 | 0.53 | 0.16 | 0.34 | -0.54 | 0.41 |
| *petB* | Q57038 | Cytochrome b6 (PetB) | -0.30 | -0.03 | 0.00 | 0.98 | -0.12 | -0.36 | -1.45 | 0.16 |
| *petC1* | P26290 | Cytochrome b6-f complex iron-sulfur subunit / Rieske (PetC1) | 0.11 | 0.76 | 0.46 | 0.61 | -0.13 | 0.66 | -0.02 | 0.35 |
| *petD* | P27589 | Cytochrome b6-f complex subunit 4 (PetD) | -0.36 | 0.55 | 0.41 | 1.43 | -0.61 | -0.92 | -0.38 | -0.46 |
| *petP* | P72798 | Cytochrome b6-f regulatory subunit PetP | -0.10 | 0.99 | 0.08 | 0.49 | -0.32 | 0.75 | 0.40 | 0.51 |

Scale Log2FC: ≥ +1.15 … 0 … ≤ -1.02

| Gene | Accession | Protein name | Log2FC at 200 µE $m^{-2}$ $s^{-1}$ (rel. to WT) | | | | Log2FC at 300 µE $m^{-2}$ $s^{-1}$ (rel. to WT) | | | |
|---|---|---|---|---|---|---|---|---|---|---|
| | | | EFE | Δglgc | Δglgc+EFE | LGS | EFE | Δglgc | Δglgc+EFE | LGS |
| **Calvin–Benson–Bassham cycle (CBB) (n=20)** | | | | | | | | | | |
| *ccmK2* | P72761 | Carboxysome shell protein CcmK2 | 0.33 | -0.84 | -0.60 | 5.26 | 0.23 | -0.09 | -0.67 | 0.41 |
| *gap2* | P80505 | Glyceraldehyde-3-phosphate dehydrogenase 2 | -0.05 | -0.33 | -0.01 | 0.27 | -4.54 | -4.42 | -5.02 | -4.51 |
| *ccmN* | P72757 | Carboxysome assembly protein CcmN | 0.07 | -1.52 | 1.08 | 0.54 | 0.55 | -0.89 | 4.79 | 0.15 |
| *cbbL* | P54205 | Ribulose bisphosphate carboxylase large chain | 4.24 | -0.76 | -1.05 | 0.27 | -0.06 | -0.51 | -1.26 | -0.04 |
| *ccmL* | P72759 | Carboxysome shell vertex protein CcmL | -0.52 | -0.93 | 0.85 | -0.41 | 0.38 | -0.51 | 3.06 | -0.23 |
| *rbcR* | P73862 | Rubisco operon transcriptional regulator | -0.84 | -0.52 | -0.76 | -1.33 | 0.18 | 0.98 | -0.28 | 2.36 |
| *ccmO* | Q55121 | Carboxysome assembly protein CcmO | -0.90 | -0.18 | 1.15 | -1.95 | 0.02 | 0.36 | 0.95 | 0.18 |
| *ccaA* | Q54735 | Carbonic anhydrase | -0.15 | -1.79 | -1.87 | -0.50 | 0.21 | 0.47 | -0.42 | 0.69 |
| *gap1* | P49433 | Glyceraldehyde-3-phosphate dehydrogenase 1 | -0.71 | -1.17 | -1.39 | -0.41 | -0.42 | -1.21 | -1.72 | -1.10 |
| *ccmK3* | P73406 | Carboxysome shell protein CcmK3 | -0.25 | -0.66 | -1.51 | 0.61 | -0.18 | -0.08 | -0.65 | -0.28 |
| *tpiA* | Q59994 | Triosephosphate isomerase | -0.16 | -1.02 | -1.33 | 0.66 | 0.21 | 0.38 | -0.61 | 0.56 |
| *ccmK4* | P73407 | Carboxysome shell protein CcmK4 | -0.14 | -0.34 | -0.53 | 0.56 | -0.59 | -0.70 | -1.26 | -0.74 |
| *ccmP* | Q55550 | Carboxysome shell protein CcmP | 0.11 | -0.17 | -0.43 | 0.52 | -0.27 | -0.75 | -1.25 | -0.30 |
| *icfA* | Q55136 | carbonic anhydrase | -0.70 | -1.11 | -1.01 | 0.17 | 0.18 | -0.32 | 0.12 | 0.33 |
| *rbcR* | P73123 | Probable RuBisCO transcriptional regulator | 0.09 | 1.08 | 0.70 | 0.22 | 0.44 | -0.09 | 0.10 | -0.36 |
| *rpiA* | Q55766 | Ribose-5-phosphate isomerase A | 0.08 | -0.48 | -0.20 | 0.00 | 0.00 | -0.08 | -1.03 | -0.21 |
| *rbcX* | Q55670 | RuBisCO chaperone RbcX | 0.54 | 0.33 | 0.76 | -1.03 | 0.33 | -0.36 | 0.62 | 0.38 |
| *ccmM* | P72758 | Carboxysome assembly protein CcmM | 0.02 | -0.54 | -0.73 | 0.25 | 0.16 | -0.39 | -1.02 | 0.35 |
| *prk* | P37101 | Phosphoribulokinase | -0.03 | -1.00 | -0.91 | 0.26 | 0.18 | -0.13 | -0.64 | 0.35 |
| *pgk* | P74421 | Phosphoglycerate kinase | -0.58 | -0.50 | -0.92 | -0.36 | -0.26 | 0.18 | -0.76 | -0.63 |
| *ccmA* | P72864 | Carboxysome formation protein | -0.05 | -0.58 | -0.44 | 0.72 | 0.40 | 0.04 | -0.81 | 0.91 |
| *ccmK1* | P72760 | Carboxysome shell protein CcmK1 | 0.28 | -0.88 | -0.71 | 0.66 | 0.12 | 0.03 | -0.76 | 0.61 |
| *fda* | P74309 | Fructose-bisphosphate aldolase class 1 | -0.08 | 0.83 | 0.28 | -0.02 | -0.16 | -0.27 | -0.17 | -0.31 |
| *ccmM* | P72758-2 | Isoform CcmM52 of Carboxysome assembly protein CcmM | 0.13 | -0.68 | -0.72 | 0.46 | 0.28 | -0.35 | -0.83 | 0.43 |
| *tktA* | P73282 | Transketolase | -0.25 | -0.80 | -0.81 | 0.14 | 0.13 | 0.37 | -0.56 | 0.22 |
| *cbbS* | P54206 | Ribulose bisphosphate carboxylase small subunit | -0.34 | -0.23 | -0.79 | 0.42 | 0.02 | 0.80 | -0.49 | 0.52 |
| *fbaA* | Q55664 | Fructose-bisphosphate aldolase class 2 | 0.24 | 0.16 | -0.04 | 0.51 | -0.03 | 0.56 | -0.05 | -0.26 |
| *fbp* | P74324 | Fructose-1,6-bisphosphatase class 1 | -0.46 | 0.49 | 0.13 | -0.23 | 0.37 | 0.13 | -0.01 | 0.52 |
| *raf1* | Q55875 | RuBisCO accumulation factor 1 | 0.40 | -0.13 | 0.05 | 0.16 | 0.09 | -0.17 | -0.13 | -0.17 |
| **Oxidative stress response (n=21)** | | | | | | | | | | |
| *gpx2* | P73824 | Hydroperoxy fatty acid reductase Gpx2 | -0.52 | -0.06 | 1.85 | -0.39 | 0.10 | 1.43 | 2.88 | 0.42 |
| *gst* | Q55139 | Glutathione S-transferase | -0.23 | 0.21 | -0.08 | -0.28 | 0.60 | -1.12 | -2.24 | -0.68 |
| *msrA2* | P72800 | Peptide methionine sulfoxide reductase MsrA 2 | 1.49 | 1.14 | 0.96 | 2.08 | -0.85 | 0.17 | -0.59 | 0.94 |
| *bcp* | P72719 | thioredoxin-dependent peroxiredoxin | 0.21 | -0.44 | 0.09 | 1.86 | -1.09 | 0.30 | 0.63 | -0.90 |
| *isiB* | P27319 | Flavodoxin | -1.35 | -1.06 | -1.80 | -0.70 | 0.19 | -0.78 | 0.57 | 0.14 |
| *slr0233* | P52232 | Thioredoxin-like protein slr0233 | 0.60 | -0.03 | 1.75 | 1.60 | 0.19 | -0.64 | -0.20 | 0.56 |
| *gshB* | P73493 | Glutathione synthetase | -0.50 | -1.41 | -1.71 | 0.26 | 0.01 | -0.84 | -1.20 | -0.41 |
| *katG* | P73911 | Catalase-peroxidase | 0.20 | 0.10 | 0.15 | 0.41 | -0.46 | -1.70 | -0.86 | -1.13 |
| *ftrC* | Q55389 | Ferredoxin-thioredoxin reductase, catalytic chain | -0.11 | 0.37 | -0.57 | -0.29 | 0.65 | 1.58 | 0.96 | 1.53 |
| *sll0755* | Q55624 | Putative peroxiredoxin sll0755 | 1.37 | -0.56 | 0.52 | 0.88 | 1.10 | 0.00 | 1.24 | 1.11 |
| *gpx1* | P74250 | Hydroperoxy fatty acid reductase gpx1 | -0.15 | -1.37 | -0.71 | 0.49 | 0.07 | 1.10 | -0.53 | -0.11 |
| *slr1963* | P74102 | Orange carotenoid-binding protein | 0.08 | -0.31 | 1.25 | -0.42 | 0.52 | -0.64 | 0.52 | -0.08 |
| *slr1139* | P73263 | Thioredoxin-like protein slr1139 | 0.48 | 0.88 | 1.02 | 0.90 | -0.10 | 1.19 | 1.08 | -0.61 |
| *ftrV* | Q55781 | Ferredoxin-thioredoxin reductase, variable chain | -0.65 | -0.62 | -1.17 | -0.74 | -0.09 | -0.03 | -0.50 | -0.01 |
| *gst1* | P74665 | Glutathione S-transferase | -0.01 | -1.14 | -0.72 | 0.26 | -0.08 | -0.32 | -1.08 | -0.19 |
| *sll1680* | P72779 | peptide-methionine (R)-S-oxide reductase | -0.65 | -0.06 | -0.56 | 1.01 | -0.06 | -0.09 | -0.86 | 0.84 |
| *trxA* | P52231 | Thioredoxin | 0.15 | 0.61 | 0.23 | 0.18 | 0.01 | 0.98 | 0.59 | 0.37 |
| *sll1621* | P73728 | Peroxiredoxin sll1621 | -0.28 | -0.41 | -0.53 | 0.14 | 0.00 | 0.72 | -0.09 | 0.90 |
| *msrA1* | P72622 | Peptide methionine sulfoxide reductase MsrA 1 | 0.08 | -0.90 | -0.67 | -0.41 | 0.10 | 0.59 | 0.07 | 0.61 |
| *bcp* | P72697 | thioredoxin-dependent peroxiredoxin | 0.00 | -0.21 | -0.55 | 0.76 | -0.01 | 0.82 | 0.37 | 0.41 |
| *sodB* | P77968 | Superoxide dismutase [Fe] | 0.20 | -0.59 | -0.03 | 0.26 | 0.54 | 0.49 | 0.40 | -0.04 |
| **Chaperones, proteases & stress proteins (n=27)** | | | | | | | | | | |
| *hliA* | P73183 | High light-inducible protein HliA | n.d. | n.d. | n.d. | n.d. | n.d. | n.d. | n.d. | n.d. |
| *clpC* | Q55662 | ATP-dependent Clp protease regulatory subunit | -0.17 | 0.62 | 0.20 | 1.16 | -0.37 | -0.42 | 3.75 | -0.60 |
| *tig* | Q55511 | Trigger factor | 0.77 | -2.29 | -1.04 | -0.25 | 0.79 | 1.31 | 1.49 | 3.62 |
| *hsp17* | P72977 | 16.6 kDa small heat shock protein, molecular chaperon | 1.82 | 2.09 | 1.48 | 0.86 | 2.28 | 0.96 | 1.71 | 0.30 |
| *clpS* | P73634 | ATP-dependent Clp protease adapter protein ClpS | -1.26 | -1.97 | -1.73 | -1.95 | -1.14 | -0.40 | -0.55 | -1.76 |
| *groES* | Q05971 | Co-chaperonin GroES | 1.65 | 0.05 | 1.86 | 0.10 | 1.19 | 0.81 | 1.83 | 0.63 |
| *clpR* | P74466 | Putative ATP-dependent Clp protease proteolytic subunit-like | 0.18 | 0.30 | 0.14 | -1.67 | 0.51 | -0.18 | 0.11 | 1.11 |
| *groEL2* | P22034 | Chaperonin GroEL 2 | 1.47 | -0.72 | 1.09 | 0.04 | 1.42 | -0.52 | 0.54 | 0.10 |
| *groEL1* | Q05972 | Chaperonin GroEL 1 | 1.42 | -0.66 | 1.19 | 0.29 | 1.12 | -0.61 | 0.27 | -0.06 |
| *hslO* | P73910 | 33 kDa chaperonin | 0.08 | 0.99 | 0.33 | 0.48 | 0.46 | 1.32 | 1.41 | 0.57 |
| *dnaK3* | P73098 | Chaperone protein dnaK3 | 1.29 | 1.27 | 0.35 | 0.01 | -0.07 | 0.11 | -0.63 | 0.00 |
| *dnaJ2* | P73097 | Chaperone protein DnaJ 2 | -0.97 | -1.16 | -0.54 | -0.56 | 0.62 | 0.38 | 1.24 | 1.06 |
| *clpP2* | Q59993 | ATP-dependent Clp protease proteolytic subunit 2 | -1.10 | -0.70 | -0.87 | -0.24 | -0.51 | 1.12 | 0.01 | -0.09 |
| *clpP3* | P74467 | Probable ATP-dependent Clp protease proteolytic subunit 3 | 0.55 | 0.22 | 0.27 | 1.02 | 0.34 | 0.40 | -0.34 | 0.12 |
| *clpB2* | P74361 | Chaperone protein ClpB 2 | -0.49 | 0.16 | -0.57 | -0.96 | 0.32 | 0.87 | -0.05 | 0.50 |
| *dnaK1* | Q55154 | Chaperone protein dnaK1 | -0.30 | -0.51 | -0.82 | -0.38 | 0.23 | 0.64 | -0.18 | 0.85 |
| *ftsH2* | Q55700 | ATP-dependent zinc metalloprotease FtsH 2 | 0.28 | 0.73 | 0.35 | 0.85 | -0.31 | -0.31 | -0.54 | -0.23 |
| *dnaJ1* | Q55505 | Chaperone protein DnaJ 1 | 0.04 | 0.13 | -0.55 | 0.43 | 0.11 | -0.19 | -0.72 | 0.33 |
| *dnaK2* | P22358 | Chaperone protein DnaK2 | 0.67 | 0.57 | 0.71 | 0.19 | 0.06 | 0.08 | 0.36 | -0.46 |
| *htpG* | P74702 | Heat shock protein | -0.23 | 0.45 | -0.03 | -0.62 | 0.24 | -0.13 | -0.17 | 0.40 |
| *ftsH4* | P73437 | ATP-dependent zinc metalloprotease FtsH 4 | -0.13 | -0.27 | -0.38 | 0.62 | 0.07 | -0.19 | -0.31 | -0.25 |
| *clpB1* | P74459 | Chaperone protein ClpB 1 | 0.61 | 0.27 | 0.03 | 0.20 | 0.19 | -0.52 | -0.29 | 0.06 |
| *clpP1* | P54416 | ATP-dependent Clp protease proteolytic subunit 1 | 0.16 | -0.21 | -0.08 | 0.56 | 0.22 | 0.25 | -0.28 | 0.43 |
| *ftsH1* | P73179 | ATP-dependent zinc metalloprotease FtsH 1 | 0.36 | -0.47 | -0.04 | 0.55 | 0.32 | -0.17 | 0.24 | -0.26 |
| *grpE* | Q59978 | Protein GrpE | 0.24 | -0.35 | -0.51 | -0.25 | 0.06 | -0.11 | -0.28 | -0.04 |
| *clpX* | Q55510 | ATP-dependent Clp protease ATP-binding subunit ClpX | 0.18 | -0.43 | 0.06 | 0.27 | 0.17 | -0.31 | -0.34 | 0.33 |
| *ftsH3* | P72991 | ATP-dependent zinc metalloprotease FtsH 3 | 0.17 | 0.26 | -0.20 | -0.40 | -0.35 | 0.00 | -0.41 | 0.27 |
| **Fatty acid synthesis (n=11)** | | | | | | | | | | |
| *acpP* | P20804 | Acyl carrier protein | 0.52 | -0.57 | 1.16 | 0.70 | 0.58 | 1.88 | 2.30 | 3.37 |
| *fabF* | P73283 | 3-oxoacyl-[acyl-carrier-protein] synthase 2 | 1.20 | 2.21 | 1.78 | 1.06 | -0.41 | -1.55 | -0.30 | -1.48 |
| *accC* | Q55160 | Biotin carboxylase | -0.13 | -2.12 | -1.19 | -0.41 | -0.18 | 0.14 | -0.40 | 0.97 |
| *accA* | P74638 | Acetyl-coenzyme A carboxylase carboxyl transferase subunit alpha | 0.49 | -0.53 | -0.34 | 1.03 | 0.14 | -0.68 | -1.83 | -0.71 |
| *fabH* | P73951 | Beta-ketoacyl-[acyl-carrier-protein] synthase III | 0.22 | 0.40 | -0.31 | -0.17 | -0.81 | -0.72 | -1.73 | -0.47 |
| *fabZ* | P73848 | 3-hydroxyacyl-[acyl-carrier-protein] dehydratase FabZ | 1.05 | 1.51 | 1.10 | 0.79 | -0.06 | -1.13 | 0.42 | -0.78 |
| *fabD* | P73242 | Malonyl CoA-acyl carrier protein transacylase | 1.33 | 1.45 | 1.30 | 1.29 | 0.04 | 0.62 | -0.23 | 0.14 |
| *fabG* | P73574 | 3-oxoacyl-[acyl-carrier-protein] reductase | -0.55 | -1.37 | -0.72 | -0.65 | -0.16 | -0.37 | 0.60 | -0.16 |
| *accD* | Q57417 | Acetyl-coenzyme A carboxylase carboxyl transferase subunit beta | -0.11 | -1.17 | -0.69 | -0.18 | 0.63 | 0.03 | -0.33 | 0.92 |
| *accB* | Q55120 | Biotin carboxyl carrier protein of acetyl-CoA carboxylase | 0.32 | -0.32 | -0.79 | 0.17 | -0.04 | -0.83 | -0.50 | -0.49 |
| *fabI* | P73016 | Enoyl-[acyl-carrier-protein] reductase [NADH] FabI | 0.22 | -0.11 | -0.03 | 0.78 | 0.16 | 0.10 | -0.74 | 0.18 |
| **Glycogen metabolism (n=11)** | | | | | | | | | | |
| *glgA2* | P72623 | Probable glycogen synthase 2 | -0.01 | -1.62 | -1.75 | 10.81 | 0.30 | -0.28 | -0.87 | -0.25 |
| *stpA* | Q55034 | Glucosylglycerol-phosphate phosphatase | -0.91 | -2.94 | -3.11 | -0.61 | 0.18 | 1.85 | -1.74 | 0.39 |
| *glgB* | P52981 | 1,4-alpha-glucan branching enzyme GlgB | -0.32 | -2.19 | -2.79 | -0.11 | 0.04 | -2.06 | -2.31 | -0.39 |
| *glgC* | P52415 | Glucose-1-phosphate adenylyltransferase | 0.40 | 0.31 | 0.50 | 0.44 | 0.15 | -2.24 | -0.55 | -0.07 |
| *glgP* | P73546 | Alpha-1,4 glucan phosphorylase | -0.29 | -1.83 | -1.82 | -0.78 | 0.45 | -0.16 | -0.58 | 0.16 |
| *glgX* | P73608 | Glycogen operon protein GlgX | 0.13 | 1.78 | 1.26 | 1.30 | -0.49 | -0.79 | -0.37 | -0.66 |
| *malQ* | P72785 | 4-alpha-glucanotransferase | -0.20 | -0.24 | -0.91 | 0.75 | 0.82 | -0.83 | 1.00 | -0.60 |
| *glgX* | P72691 | Glycogen operon protein GlgX | 0.63 | -0.88 | -0.66 | 0.58 | 0.14 | -0.81 | -0.98 | -0.39 |
| *glgA1* | P74521 | Glycogen synthase 1 | 0.14 | -0.34 | -0.59 | 0.21 | -0.03 | -0.05 | -0.39 | -0.18 |
| *glgP* | P73511 | Glycogen phosphorylase | 0.21 | -0.11 | -0.18 | 0.08 | 0.33 | -0.53 | 0.21 | 0.28 |
| *ggpS* | P74258 | Glucosylglycerol-phosphate synthase | -0.03 | -0.35 | 0.15 | 0.08 | 0.05 | -0.11 | 0.17 | 0.06 |
| **Sucrose metabolism (n=2)** | | | | | | | | | | |
| *slr0953* | P74325 | sucrose-phosphate phosphatase | 0.50 | -1.47 | -0.73 | -2.45 | 0.68 | 1.46 | -1.17 | -1.08 |
| *sps* | Q55440 | sucrose-phosphate synthase | | | | | | | | |
| **Polyhydroxybutyrate (PHB) (n=5)** | | | | | | | | | | |
| *phaP* | P73545 | Phasin PhaP | 0.29 | -0.68 | -0.45 | -0.13 | -0.19 | -2.61 | -1.48 | -1.52 |
| *phaA* | P73825 | Acetyl-CoA acetyltransferase | 0.09 | 0.99 | 1.98 | -1.58 | 0.72 | 0.01 | 2.34 | 2.33 |
| *phaE* | P73389 | Poly(3-hydroxyalkanoate) polymerase subunit PhaE | 1.28 | -1.06 | 1.07 | -0.51 | 0.24 | -2.00 | -1.37 | -1.76 |
| *phaB* | P73826 | Acetoacetyl-CoA reductase | -0.28 | 0.34 | 0.25 | -0.79 | -0.28 | -0.69 | -0.13 | -0.41 |
| *phaC* | P73390 | Poly(3-hydroxyalkanoate) polymerase subunit PhaC | -0.45 | 0.65 | -0.05 | 0.27 | -0.04 | 0.20 | -0.05 | -0.17 |

| Gene | Accession | Protein name | Log2FC at 200 µE $m^{-2}$ $s^{-1}$ (rel. to WT) | | | | Log2FC at 300 µE $m^{-2}$ $s^{-1}$ (rel. to WT) | | | |
|---|---|---|---|---|---|---|---|---|---|---|
| | | | EFE | Δglgc | Δglgc+EFE | LGS | EFE | Δglgc | Δglgc+EFE | LGS |
| **Glycolysis (n=15)** | | | | | | | | | | |
| *pfkA2* | Q55988 | ATP-dependent 6-phosphofructokinase 2 | -0.41 | -0.67 | 0.21 | 0.35 | -17.77 | -17.64 | -18.18 | -16.81 |
| *cscK* | P73521 | Fructokinase | 0.01 | 0.07 | -2.74 | 0.27 | 0.21 | 0.24 | 1.53 | 0.09 |
| *glk* | Q55855 | Glucokinase | -0.99 | -2.07 | -1.22 | 0.76 | -0.05 | -0.17 | -0.55 | -1.15 |
| *pdhA* | P74490 | Pyruvate dehydrogenase E1 component subunit alpha | -0.23 | -1.18 | -1.36 | 0.93 | -0.21 | -0.64 | -1.13 | -0.25 |
| *pdhB* | P73405 | Pyruvate dehydrogenase E1 beta subunit | -0.30 | -0.47 | -0.07 | -0.79 | 0.17 | 0.51 | 0.33 | 1.29 |
| *sll0395* | Q55734 | phosphoglycerate mutase (2,3-diphosphoglycerate-dependent) | 0.94 | -0.76 | -1.23 | -0.07 | 0.25 | 0.32 | -0.49 | 1.04 |
| *gnd* | P52208 | 6-phosphogluconate dehydrogenase, decarboxylating | -0.39 | -0.95 | -1.02 | -0.28 | -0.43 | -1.10 | -0.99 | -1.20 |
| *pyk1* | Q55863 | Pyruvate kinase 1 | -0.07 | 0.16 | 0.13 | 0.13 | 0.38 | -0.49 | 0.01 | -1.11 |
| *gpmB* | P72649 | Phosphoglycerate mutase | -0.72 | -0.40 | -1.10 | -0.68 | 0.40 | 0.55 | 0.07 | 0.91 |
| *eno* | P77972 | Enolase | -0.31 | -0.32 | -0.55 | 0.46 | -0.16 | -0.23 | -1.02 | 0.08 |
| *pfkA1* | P72830 | ATP-dependent 6-phosphofructokinase 1 | 0.08 | -0.15 | -0.49 | 0.24 | 0.21 | 0.13 | -0.16 | 1.00 |
| *gpmI* | P74507 | 2,3-bisphosphoglycerate-independent phosphoglycerate mutase | 0.07 | -0.12 | -0.39 | 0.01 | 0.10 | -0.09 | -0.18 | 0.81 |
| *pyk2* | P73534 | Pyruvate kinase 2 | 0.15 | -0.68 | -0.09 | 0.48 | 0.56 | 0.32 | -0.20 | 0.47 |
| *pgm* | P74643 | Phosphoglucomutase | 0.21 | 0.68 | 0.15 | 0.53 | -0.28 | 0.25 | -0.52 | -0.12 |
| *pgi* | P52983 | Glucose-6-phosphate isomerase | 0.19 | -0.03 | -0.27 | -0.31 | 0.32 | 0.05 | -0.38 | -0.09 |
| **TCA cycle (n=10)** | | | | | | | | | | |
| *mdh* | Q55383 | Malate dehydrogenase | -0.09 | -1.00 | -0.74 | -0.22 | 0.28 | 2.65 | -0.73 | 0.62 |
| *icd* | P80046 | Isocitrate dehydrogenase [NADP] | -0.22 | -1.33 | -1.33 | -0.13 | 0.05 | 0.71 | -0.82 | 0.39 |
| *gltA* | Q59977 | Citrate synthase | 0.18 | -1.06 | -1.06 | 0.45 | 0.27 | -0.52 | -0.30 | 0.38 |
| *sucC* | P72927 | Succinate--CoA ligase | 0.85 | 0.37 | 0.78 | 1.06 | 0.67 | 0.13 | 0.35 | 0.28 |
| *sucD* | P74286 | Succinyl-CoA synthetase | | | | | 0.97 | | 0.24 | |
| *fumC* | Q55674 | Fumarate hydratase class II | 0.23 | -0.37 | -0.89 | -0.24 | -0.22 | -0.79 | -0.47 | -0.23 |
| *odhB* | P74510 | Dihydrolipoamide acetyltransferase component of pyruvate dehydrogenase complex | 0.03 | -0.66 | -0.21 | -0.31 | 0.66 | 0.36 | -0.34 | 0.23 |
| *ppc* | P74299 | Phosphoenolpyruvate carboxylase | 0.61 | 0.52 | 0.34 | 0.24 | 0.46 | 0.60 | 0.17 | 0.07 |
| *acnB* | P74582 | Aconitate hydratase B | -0.03 | 0.06 | 0.43 | 0.30 | 0.16 | -0.04 | -0.55 | 0.41 |
| *me* | P72661 | Malic enzyme | -0.39 | 0.09 | -0.26 | 0.24 | -0.02 | 0.12 | -0.19 | 0.20 |
| **Pentose phosphate pathway (PPP) (n=8)** | | | | | | | | | | |
| *pgl* | P74618 | 6-phosphogluconolactonase | -0.96 | -1.70 | -2.45 | -0.60 | -0.07 | 0.03 | -2.00 | -0.48 |
| *gnd* | P52208 | 6-phosphogluconate dehydrogenase, decarboxylating | -0.39 | -0.95 | -1.02 | -0.28 | -0.43 | -1.10 | -0.99 | -1.20 |
| *zwf* | P73411 | Glucose-6-phosphate 1-dehydrogenase | -0.35 | -1.14 | -1.15 | 0.08 | 0.02 | -0.87 | -1.14 | -0.61 |
| *rpiA* | Q55766 | Ribose-5-phosphate isomerase A | 0.08 | -0.48 | -0.20 | 0.00 | 0.00 | -0.08 | -1.03 | -0.21 |
| *tktA* | P73282 | Transketolase | -0.25 | -0.80 | -0.81 | 0.14 | 0.13 | 0.37 | -0.56 | 0.22 |
| *slr2094* | P73922 | D-fructose 1,6-bisphosphatase class 2/sedoheptulose 1,7-bisphosphatase | 0.15 | 0.64 | 0.28 | -0.50 | 0.26 | 0.19 | 0.34 | 0.35 |
| *tal* | P72797 | Transaldolase | 0.05 | 0.10 | -0.12 | 0.22 | 0.01 | -0.15 | -0.59 | -0.61 |
| *rpe* | P74061 | Ribulose-phosphate 3-epimerase | 0.20 | -0.33 | -0.35 | 0.27 | 0.23 | 0.18 | 0.08 | 0.06 |
| **Nitrogen metabolism (n=16)** | | | | | | | | | | |
| *gdhA* | P54386 | NADP-specific glutamate dehydrogenase | -0.13 | -1.13 | -1.21 | 8.81 | 0.04 | 1.80 | -0.77 | 0.61 |
| *sll1017* | P72935 | Putative ammonium transporter sll1017 | 1.79 | | 6.97 | 8.77 | | | | |
| *nrtD* | P73265 | Nitrate import ATP-binding protein NrtD | 0.09 | -1.98 | -3.68 | 0.30 | 1.30 | 0.99 | 0.33 | 0.28 |
| *narB* | P73448 | Nitrate reductase | -0.09 | -1.13 | 1.16 | -1.06 | 0.39 | -0.19 | 1.89 | 0.02 |
| *nrtB* | P73451 | Nitrate import permease protein NrtB | 0.66 | -0.87 | -0.42 | 1.33 | 0.00 | -0.44 | -1.46 | -0.24 |
| *glnA* | P77961 | Glutamine synthetase | -0.40 | -0.82 | -1.43 | -0.37 | -0.12 | 0.30 | -1.07 | -0.15 |
| *glnN* | P77970 | Glutamate--ammonia ligase | -0.63 | -0.77 | -1.00 | 0.57 | -0.42 | -0.13 | -1.15 | 0.66 |
| *nrtC* | P73450 | Nitrate import ATP-binding protein NrtC | -0.15 | -0.93 | -0.80 | 0.37 | -0.18 | -0.86 | -1.14 | 0.28 |
| *nrtD* | P73449 | Nitrate transport protein NrtD | 0.15 | -0.09 | -0.38 | 0.68 | 0.56 | 0.38 | -0.96 | 0.88 |
| *gltD* | P72762 | NADH-glutamate synthase small subunit | 0.61 | -0.30 | 0.77 | 0.10 | 0.65 | -0.25 | 0.94 | 0.00 |
| *sll0108* | P54147 | Putative ammonium transporter sll0108 | -0.02 | 0.45 | 0.26 | 0.06 | -0.32 | -0.89 | -0.67 | 0.31 |
| *nrtA* | P73452 | Nitrate/nitrite binding protein NrtA | 0.12 | -0.50 | -0.42 | 0.33 | 0.09 | -0.18 | -0.75 | 0.23 |
| *gltB* | P55037 | Ferredoxin-dependent glutamate synthase 1 | 0.13 | 0.24 | -0.16 | -0.65 | 0.26 | -0.25 | 0.28 | 0.05 |
| *gltS* | P55038 | Ferredoxin-dependent glutamate synthase 2 | 0.14 | -0.61 | -0.36 | 0.42 | 0.21 | -0.32 | -0.63 | -0.02 |
| *nirA* | Q55366 | Ferredoxin--nitrite reductase | -0.09 | -0.25 | -0.19 | 0.50 | 0.14 | -0.46 | -0.60 | 0.06 |
| *glnB* | Q55247 | Nitrogen regulatory protein P-II | 0.20 | 0.06 | -0.17 | 0.34 | -0.21 | 0.29 | -0.16 | -0.31 |
| **Cyanophycin (n=2)** | | | | | | | | | | |
| *cphB* | P73832 | Cyanophycinase | 0.03 | -0.78 | -0.71 | 0.01 | 0.28 | -0.60 | -0.35 | -0.27 |
| *cphA* | P73833 | Cyanophycin synthetase | -0.19 | -0.57 | -0.76 | 0.17 | -0.14 | -0.04 | -0.64 | 0.07 |
| **Polyphosphate (n=2)** | | | | | | | | | | |
| *ppk* | Q55898 | Polyphosphate kinase | 0.10 | 0.40 | 0.30 | 0.32 | -0.04 | 0.24 | -0.11 | 0.32 |
| *ppx* | P74663 | Exopolyphosphatase | -0.08 | 0.22 | 0.04 | -0.54 | 0.72 | 1.36 | 1.43 | 1.57 |
| **Phosphate Transport (n=6)** | | | | | | | | | | |
| *pstC* | Q55198 | Phosphate transport system permease protein | 0.18 | | 2.97 | -5.38 | | -0.11 | 8.37 | 0.46 |
| *pstS* | P73785 | Phosphate-binding protein | -0.70 | 0.11 | 2.22 | -0.65 | 0.16 | -0.23 | 5.10 | -0.58 |
| *pstC* | P73786 | Phosphate transport system permease protein | -4.56 | -0.90 | 0.40 | -0.84 | 0.30 | -2.11 | 0.79 | -1.27 |
| *pstS* | Q55199 | Phosphate-binding protein | 0.30 | -0.49 | 1.25 | 0.05 | 0.44 | 1.73 | 3.20 | -0.14 |
| *pstB1* | Q55196 | Phosphate import ATP-binding protein PstB 1 | -1.14 | -1.03 | 2.12 | -1.22 | 0.18 | -0.17 | 1.81 | -0.93 |
| *pstB2* | Q55195 | Phosphate import ATP-binding protein PstB 2 | -0.46 | -1.55 | -0.17 | -1.19 | 0.34 | -0.06 | 0.93 | 0.73 |
| **Phosphate Signaling (n=2)** | | | | | | | | | | |
| *sphX* | Q55200 | Protein SphX | -0.49 | -0.67 | 2.10 | -0.46 | -0.22 | -0.36 | 2.45 | 0.13 |
| *slr1983* | P74139 | Histidine kinase SphS (slr1983) | -0.18 | -1.85 | 0.41 | -0.37 | 0.63 | -0.43 | 2.81 | 0.74 |
| **Iron & Cation Transport (n=10)** | | | | | | | | | | |
| *slr0794* | Q55935 | Cation or drug efflux system protein | -0.66 | 0.13 | 1.91 | 0.09 | 0.20 | 0.59 | 4.57 | 0.20 |
| *ziaA* | Q59998 | Zinc-transporting ATPase | -1.43 | -2.31 | -1.44 | -2.03 | 1.68 | 1.01 | 2.61 | 2.01 |
| *slr0454* | P74693 | Cation or drug efflux system protein | -0.53 | -1.29 | -2.03 | 0.27 | -0.10 | -1.49 | -2.48 | 0.47 |
| *pacS* | P73241 | Probable copper-transporting ATPase PacS | 0.75 | 0.13 | -0.03 | -0.43 | -0.09 | -2.09 | -0.14 | -1.59 |
| *kdpB* | P73867 | Potassium-transporting ATPase ATP-binding subunit | 0.22 | 0.57 | -0.04 | 0.45 | -0.32 | -1.83 | -0.71 | -1.75 |
| *futA2* | Q55835 | Iron uptake protein A2 | 0.03 | -0.13 | -1.02 | 0.21 | -0.40 | -1.34 | -1.34 | -0.12 |
| *futA1* | P72827 | Iron uptake protein A1 | -0.54 | 0.09 | -0.75 | 0.25 | -0.57 | -0.08 | -1.18 | 0.17 |
| *slr2131* | P73998 | Cation or drug efflux system protein | 0.40 | 0.35 | 0.30 | -0.01 | 0.55 | 0.45 | 0.71 | 1.08 |
| *kdpA* | P73866 | Potassium-transporting ATPase potassium-binding subunit | -0.52 | -0.19 | -0.86 | 0.40 | -0.50 | -0.19 | -1.04 | -0.52 |
| *slr0369* | Q55584 | Cation or drug efflux system protein | -0.09 | -0.76 | -0.47 | 0.34 | 0.12 | -0.39 | -0.65 | 0.31 |
| **Amino Acid Transport (n=4)** | | | | | | | | | | |
| *livF* | P73650 | High-affinity branched-chain amino acid transport ATP-binding protein BraG | 7.38 | 1.12 | 7.35 | 0.53 | 7.21 | -0.28 | 7.45 | 0.46 |
| *braG* | Q55753 | High-affinity branched-chain amino acid transport ATP-binding protein | -0.75 | -1.25 | -2.77 | -1.13 | -0.39 | -0.77 | -2.11 | -0.07 |
| *glnH* | P74223 | Glutamine-binding protein | 0.36 | -1.14 | -1.20 | 0.04 | -0.04 | -2.10 | -1.31 | -0.44 |
| *livG* | Q55614 | High-affinity branched-chain amino acid transport ATP-binding protein | 0.76 | 1.64 | 1.07 | 0.74 | -0.14 | 0.38 | 0.08 | -0.09 |
| **Two-Component Signaling (n=14)** | | | | | | | | | | |
| *sll0038* | Q55448 | PatA subfamily response regulator | -0.25 | -0.88 | 1.59 | 0.13 | 0.22 | -0.48 | 4.57 | 0.12 |
| *slr1042* | P73006 | CheY subfamily response regulator | 3.51 | -1.06 | 3.03 | 0.61 | 2.96 | -0.26 | 2.53 | -0.36 |
| *sll1330* | P74541 | OmpR subfamily response regulator | -0.61 | 0.14 | 0.34 | 1.95 | -0.52 | 0.63 | -0.11 | 2.97 |
| *slr1983* | P74139 | Histidine kinase | -0.18 | -1.85 | 0.41 | -0.37 | 0.63 | -0.43 | 2.81 | 0.74 |
| *slr0947* | P74314 | Probable transcriptional regulator Ycf27 | -0.01 | -1.27 | -1.79 | 0.89 | -0.19 | -0.61 | -2.46 | -0.23 |
| *sll1003* | P73025 | Histidine kinase | -0.73 | -2.26 | -1.76 | -1.10 | 0.93 | 0.62 | 1.13 | 2.11 |
| *sll5060* | Q6ZES0 | Circadian input-output histidine kinase CikA | 0.63 | 0.96 | -0.03 | 1.44 | -0.73 | -2.16 | -1.57 | -1.80 |
| *slr2104* | P73932 | Circadian input-output histidine kinase CikA | 0.69 | -0.21 | -0.71 | 2.04 | -0.22 | 0.04 | 2.15 | 0.60 |
| *ETR1* | P73363 | Histidine kinase | -0.91 | -1.86 | -2.02 | 0.43 | 0.58 | 0.09 | -1.20 | 1.56 |
| *slr0210* | Q55783 | Histidine kinase | 1.26 | 0.68 | 1.73 | 0.12 | -0.44 | -1.62 | 0.31 | -1.77 |
| *sll0474* | Q55838 | Histidine kinase (Hik28) | 0.92 | 1.62 | 1.52 | 1.61 | -0.56 | -0.59 | -0.78 | -0.94 |
| *sll1590* | P73865 | Histidine kinase | -1.02 | -0.61 | -1.43 | -0.37 | 0.07 | -0.68 | -0.71 | -1.53 |
| *sll1871* | P74137 | Circadian input-output histidine kinase CikA | 0.70 | -0.37 | 0.17 | 0.88 | -0.51 | -1.49 | -0.89 | -1.22 |
| *pilJ* | P73173 | PilJ protein | 0.29 | 0.44 | 1.35 | -0.17 | -0.40 | -0.92 | 0.91 | -0.33 |
| **Amino Acid Biosynthesis (n=10)** | | | | | | | | | | |
| *trpE2* | P74130 | Anthranilate synthase component I-like protein | -1.19 | -0.78 | 1.97 | -3.89 | 0.57 | 0.31 | 5.94 | 2.53 |
| *ilvH* | Q55141 | Acetolactate synthase small subunit | -0.14 | -1.91 | 1.49 | -0.20 | 0.37 | 0.15 | 4.90 | 0.07 |
| *ilvB* | P73918 | Acetolactate synthase | 0.89 | 1.30 | 1.55 | -0.11 | 0.09 | -1.20 | 0.89 | -0.22 |
| *thrC* | P74193 | Threonine synthase | -0.26 | -0.64 | -0.75 | 0.28 | 0.14 | 1.44 | 0.10 | 0.64 |
| *aroC* | P23353 | Chorismate synthase | -0.23 | 1.19 | 0.52 | 0.40 | -0.74 | -0.52 | -0.81 | -1.08 |
| *ilvC* | P29107 | Ketol-acid reductoisomerase (NADP+) | -0.09 | -0.32 | -0.80 | 0.31 | -0.07 | -0.04 | -1.15 | 0.29 |
| *trpE* | P20170 | Anthranilate synthase component 1 | -0.20 | -1.06 | -0.85 | 0.12 | 0.38 | -0.15 | -0.72 | 0.55 |
| *trpA* | P77960 | Tryptophan synthase alpha chain | -0.45 | -0.85 | -0.84 | 0.20 | 0.34 | -0.23 | -1.04 | -0.09 |
| *argC* | P54899 | N-acetyl-gamma-glutamyl-phosphate reductase | 0.14 | -0.66 | -0.98 | 0.12 | -0.46 | -0.48 | -0.89 | -0.61 |
| *thrC* | P73711 | Threonine synthase | 0.46 | -0.37 | -0.14 | -0.25 | 0.16 | -0.65 | -0.34 | 0.05 |
| **Chlorophyll / Tetrapyrrole Biosynthesis (n=10)** | | | | | | | | | | |
| *chlL* | P28373 | Light-independent protochlorophyllide reductase subunit L | 0.14 | -3.01 | -2.96 | -0.31 | 0.34 | -5.14 | -2.88 | -0.09 |
| *chlN* | P28372 | Light-independent protochlorophyllide reductase subunit N | -0.34 | -3.26 | -3.75 | 1.05 | -0.20 | -0.55 | -2.64 | -0.30 |
| *hemJ* | P72793 | Protoporphyrinogen IX oxidase | -0.40 | -0.02 | -0.59 | 0.42 | -0.71 | -0.91 | -1.57 | -0.85 |
| *por* | Q59987 | Light-dependent protochlorophyllide reductase (LPOR) | -0.66 | -0.59 | -0.86 | 0.03 | -0.51 | -0.15 | -1.42 | -0.39 |
| *cobA* | P73874 | Cob(I)alamin adenosyltransferase | 0.09 | -0.93 | -1.01 | 1.42 | 0.25 | -0.60 | -0.86 | -0.45 |
| *hemE* | P54224 | Uroporphyrinogen decarboxylase | -0.19 | -1.04 | -1.38 | -0.01 | -0.04 | -0.09 | -0.81 | 0.12 |
| *hemF* | P72848 | Oxygen-dependent coproporphyrinogen-III oxidase | 0.16 | -0.68 | -0.53 | 0.48 | -0.17 | -1.07 | -1.24 | -0.28 |
| *acsF1* | P72584 | Magnesium-protoporphyrin IX monomethyl ester cyclase 1 | -0.43 | -0.61 | -0.59 | 0.04 | -0.24 | -0.25 | -1.17 | 0.16 |
| *chlB* | Q55607 | Light-independent protochlorophyllide reductase subunit B | 0.01 | -0.50 | -0.66 | 0.02 | 0.12 | -1.17 | -0.67 | -0.35 |
| *cobA* | Q55749 | Uroporphyrinogen-III C-methyltransferase | | | | | | | | |

| Gene | Accession | Protein name | Log2FC at 200 µE m⁻² s⁻¹ (rel. to WT) | | | | Log2FC at 300 µE m⁻² s⁻¹ (rel. to WT) | | | |
|---|---|---|---|---|---|---|---|---|---|---|
| | | | EFE | Δglgc | Δglgc+EFE | LGS | EFE | Δglgc | Δglgc+EFE | LGS |
| **DNA Repair & Replication (n=11)** | | | | | | | | | | |
| *gyrA* | Q55738 | DNA gyrase subunit A | -0.09 | 5.68 | 5.38 | 5.12 | 0.10 | 3.91 | 4.61 | 4.72 |
| *mutT* | P72658 | 8-oxo-dGTP diphosphatase | 1.40 | -2.46 | -0.44 | 1.00 | 1.74 | 3.82 | 0.67 | 1.91 |
| *recA* | P74737 | Protein RecA | -0.41 | 0.34 | 2.47 | -1.53 | 0.56 | 0.17 | 3.80 | 0.78 |
| *recJ* | P73518 | Single-stranded DNA-specific exonuclease RecJ | -0.66 | -3.49 | -3.47 | 0.44 | 0.44 | 0.30 | 0.10 | 1.37 |
| *ruvA* | P73554 | Holliday junction branch migration complex subunit RuvA | 3.09 | -0.46 | 2.30 | 2.45 | 0.94 | 0.07 | -0.01 | -1.33 |
| *rnhA* | Q55801 | Ribonuclease HI | -0.48 | -2.77 | -0.80 | 0.28 | 0.23 | -0.09 | -0.85 | 0.15 |
| *sll1366* | P74552 | Helicase of the SNF2/RAD54 family | -0.28 | -1.68 | -2.09 | 0.53 | -0.34 | -0.10 | -0.79 | 0.37 |
| *uvrC* | P73580 | UvrABC system protein C | 0.71 | 0.13 | 0.05 | -0.49 | -0.02 | -2.05 | -0.09 | -1.35 |
| *uvrA* | P73412 | UvrABC system protein A | -0.75 | -0.52 | -1.04 | -0.45 | 0.22 | -0.36 | -1.19 | -0.28 |
| *nrdA* | P74240 | Ribonucleoside-diphosphate reductase subunit alpha | 0.60 | -0.32 | -0.19 | 0.68 | 0.53 | 0.28 | -0.48 | 1.07 |
| *hup* | P73418 | DNA-binding protein HU | 0.30 | 0.71 | 0.65 | 0.89 | -0.27 | -0.10 | -0.18 | -0.13 |
| **Proteosynthesis (n=103)** | | | | | | | | | | |
| *rpmI* | P48959 | Large ribosomal subunit protein bL35 | -1.06 | -2.17 | -2.59 | -1.00 | 0.89 | 5.50 | -2.53 | 0.91 |
| *rplF* | P73306 | Large ribosomal subunit protein uL6 | 0.96 | 1.05 | 1.40 | 5.09 | -0.14 | -0.81 | 0.70 | -0.46 |
| *rpmH* | Q55004 | Large ribosomal subunit protein bL34 | -0.58 | 0.20 | 1.29 | 0.50 | 1.08 | 1.38 | 0.55 | 4.60 |
| *ileS* | P73505 | Isoleucine–tRNA ligase | -2.38 | -2.51 | -0.88 | -4.16 | 0.21 | -0.81 | 2.73 | 2.29 |
| *rsgA* | P52640 | Small ribosomal subunit biogenesis GTPase RsgA | 0.37 | -0.37 | -3.53 | -0.13 | | | | |
| *leuS* | P73274 | Leucine–tRNA ligase | 0.37 | 0.95 | 0.81 | 0.91 | -1.40 | -1.38 | -1.22 | 3.36 |
| *rpsB* | P74071 | Small ribosomal subunit protein uS2 | 0.29 | -1.00 | 0.15 | 0.49 | 1.01 | 2.93 | 2.58 | 2.70 |
| *rpmJ* | P73300 | Large ribosomal subunit protein bL36 | -1.01 | -0.74 | -2.88 | 0.54 | 0.77 | 0.73 | -0.96 | 1.69 |
| *rpsD* | P48939 | Small ribosomal subunit protein uS4 | 0.46 | -0.20 | -0.12 | 1.07 | 0.13 | -0.55 | -2.19 | -0.10 |
| *rpmG* | P48958 | Large ribosomal subunit protein bL33 | -0.93 | -1.92 | -2.18 | -0.18 | 0.46 | 0.72 | -0.36 | 1.20 |
| *rimM* | P74035 | Ribosome maturation factor RimM | 2.12 | 0.37 | 1.65 | 1.86 | 1.57 | 0.51 | 0.94 | 1.11 |
| *alaS* | P74423 | Alanine–tRNA ligase | 0.30 | -0.23 | -0.67 | 0.36 | 0.59 | 1.99 | 0.00 | 0.61 |
| *rpsL* | P74230 | Small ribosomal subunit protein uS12 | 0.43 | 0.37 | -0.05 | 1.23 | -0.35 | -0.12 | -1.89 | 0.23 |
| *rplK* | P36237 | Large ribosomal subunit protein uL11 | -0.32 | -1.83 | -1.42 | 0.39 | 0.21 | -0.24 | -0.99 | -0.11 |
| *prfC* | P73473 | Peptide chain release factor 3 | 0.56 | 0.27 | 0.55 | 0.56 | 0.31 | 1.80 | -0.02 | 0.53 |
| *hflX* | P73965 | GTPase HflX | -1.38 | -1.71 | 0.14 | -0.12 | 0.39 | -0.32 | 0.37 | 1.18 |
| *metG* | Q55729 | Methionine–tRNA ligase | -0.14 | -1.12 | -0.86 | 0.37 | 0.58 | 1.70 | 0.31 | 1.07 |
| *hpf* | P74518 | Ribosome hibernation promotion factor | -0.09 | -1.38 | -1.69 | 0.70 | 0.28 | 0.14 | -0.99 | 0.61 |
| *rplB* | P73317 | Large ribosomal subunit protein uL2 | 0.05 | -0.59 | -0.42 | 0.29 | -0.14 | -0.58 | -1.65 | -0.02 |
| *rimP* | P72687 | Ribosome maturation factor RimP | 0.77 | 0.93 | 0.26 | 0.24 | 0.77 | 1.59 | -0.07 | 1.20 |
| *def* | P73441 | Peptide deformylase | 0.60 | 0.44 | 0.70 | 1.59 | 0.20 | 0.81 | 0.04 | 0.97 |
| *slr0923* | Q55385 | Probable small ribosomal subunit protein cS23 | 0.38 | 0.72 | 0.61 | 0.53 | -0.69 | -1.15 | -0.37 | -1.57 |
| *rplY* | P73289 | Large ribosomal subunit protein bL25 | -0.23 | -1.28 | -1.54 | 0.61 | 0.20 | 0.75 | -0.58 | 0.80 |
| *glyS* | Q55690 | Glycine–tRNA ligase beta subunit | 0.41 | -0.02 | 1.00 | 0.50 | 1.48 | -0.14 | 0.94 | -0.36 |
| *rbfA* | Q55625 | Ribosome-binding factor A | 0.77 | 0.42 | -0.25 | 0.90 | 0.66 | 1.48 | 0.84 | 0.71 |
| *rpmF* | P73014 | Large ribosomal subunit protein bL32 | 0.20 | 1.15 | 0.85 | -0.02 | 0.40 | 1.18 | 1.34 | 1.37 |
| *rplX* | P73309 | Large ribosomal subunit protein uL24 | 0.49 | -0.11 | -0.30 | 0.25 | -0.13 | -0.37 | -1.35 | -0.01 |
| *rps1A* | P73530 | Small ribosomal subunit protein bS1A | 0.36 | 0.79 | 0.36 | 0.57 | 0.32 | 1.34 | 0.98 | 0.64 |
| *proS* | P73942 | Proline–tRNA ligase | -0.09 | -1.31 | -1.33 | 0.13 | 0.24 | -0.01 | -0.26 | 0.51 |
| *prfA* | P74707 | Peptide chain release factor 1 | 0.54 | 0.20 | 0.20 | -0.03 | 0.12 | 1.31 | 0.51 | 0.40 |
| *rplU* | P74266 | Large ribosomal subunit protein bL21 | -0.29 | -1.05 | -1.31 | 0.28 | 0.75 | 0.70 | -0.66 | 0.98 |
| *argS* | Q55486 | Arginine–tRNA ligase | 0.25 | 1.29 | 0.57 | 1.10 | -0.35 | -1.00 | -0.48 | -0.98 |
| *rplT* | P48957 | Large ribosomal subunit protein bL20 | 0.92 | 0.69 | 0.86 | 1.27 | -0.26 | -0.41 | -0.80 | 0.29 |
| *frr* | P74456 | Ribosome-recycling factor | 0.31 | 0.99 | 0.59 | 0.63 | 0.03 | 1.21 | 0.53 | 0.63 |
| *gltX* | Q55778 | Glutamate–tRNA ligase | 0.19 | -0.57 | -0.40 | 0.47 | 0.46 | -0.11 | -1.20 | 0.39 |
| *rplE* | P73308 | Large ribosomal subunit protein uL5 | 0.13 | -0.67 | -0.51 | 0.54 | 0.01 | -0.42 | -1.19 | -0.27 |
| *rpsF* | P73636 | Small ribosomal subunit protein bS6 | 0.02 | 0.27 | 0.36 | 0.25 | 0.41 | 1.18 | 0.19 | 1.08 |
| *trpS* | P73655 | Tryptophan–tRNA ligase | 0.18 | -0.80 | 0.16 | 0.10 | 0.23 | -1.15 | -1.17 | -0.30 |
| *rplM* | P73294 | Large ribosomal subunit protein uL13 | 0.06 | -1.17 | -1.08 | 0.65 | 0.27 | -0.01 | -0.78 | 0.16 |
| *rpmE* | P73292 | Large ribosomal subunit protein bL31 | 0.28 | 0.55 | 0.34 | 0.73 | 0.25 | 1.07 | 0.33 | 1.17 |
| *rpsI* | P73293 | Small ribosomal subunit protein uS9 | 0.29 | -0.60 | -0.64 | 0.86 | 0.05 | -0.46 | -1.17 | -0.12 |
| *infA* | P73301 | Translation initiation factor IF-1 | -0.18 | -0.75 | -0.70 | 0.61 | 0.34 | 1.16 | 0.30 | 0.87 |
| *rplL* | P23349 | Large ribosomal subunit protein bL12 | 0.47 | 0.62 | 0.48 | 0.19 | -0.14 | -0.78 | -0.18 | -1.15 |
| *era* | Q55526 | GTPase Era | -0.20 | -0.23 | 0.00 | 0.38 | 1.15 | 0.68 | -0.45 | 0.78 |
| *rplR* | P73305 | Large ribosomal subunit protein uL18 | 0.25 | -0.14 | -0.06 | 0.87 | 0.10 | -0.65 | -1.14 | -0.08 |
| *tyrS* | P73141 | Tyrosine–tRNA ligase | -1.00 | -0.65 | -0.48 | -0.19 | 0.05 | -1.13 | -0.91 | -0.21 |
| *obg* | P72931 | GTPase Obg | -0.35 | 0.26 | 0.55 | 0.01 | -0.45 | -0.67 | -1.12 | -0.95 |
| *rpsU* | P48949 | Small ribosomal subunit protein bS21 | -0.02 | -0.37 | -0.58 | 0.87 | 0.09 | 0.72 | -0.86 | 1.10 |
| *lysS* | P73443 | Lysine–tRNA ligase | 0.39 | 0.01 | -0.28 | 1.09 | 0.18 | -0.57 | -1.03 | -0.07 |
| *rplO* | P73303 | Large ribosomal subunit protein uL15 | -0.07 | -0.47 | -0.49 | 0.05 | 0.31 | -0.27 | -1.07 | -0.11 |
| *glyQ* | Q55716 | Glycine–tRNA ligase alpha subunit | -0.25 | -1.05 | -0.81 | 0.20 | 0.15 | -0.13 | -0.94 | 0.18 |
| *prfB* | P74476 | Peptide chain release factor 2 | 0.07 | 1.04 | 0.79 | -0.02 | -0.06 | 0.35 | 0.37 | -0.25 |
| *rplI* | P42352 | Large ribosomal subunit protein bL9 | -0.06 | -0.83 | -0.79 | 0.37 | 0.02 | -0.34 | -1.02 | -0.18 |
| *rplC* | P73320 | Large ribosomal subunit protein uL3 | 0.17 | -0.53 | -0.78 | 0.36 | 0.17 | -0.05 | -1.02 | 0.46 |
| *rplQ* | P73296 | Large ribosomal subunit protein bL17 | 0.27 | -0.28 | -1.02 | 0.30 | 0.13 | -0.10 | -0.07 | 0.98 |
| *rpsR* | P48946 | Small ribosomal subunit protein bS18 | -0.15 | 1.02 | 0.24 | 0.29 | -0.39 | -0.04 | -0.55 | -0.23 |
| *rplW* | P73318 | Large ribosomal subunit protein uL23 | -0.49 | -0.43 | -0.54 | 0.70 | 0.27 | -0.32 | -1.01 | -0.40 |
| *rplV* | P73315 | Large ribosomal subunit protein uL22 | 0.09 | -0.53 | -0.52 | 0.84 | 0.17 | 0.07 | -1.01 | 0.45 |
| *rpsM* | P73299 | Small ribosomal subunit protein uS13 | 0.57 | -0.46 | 0.02 | 0.58 | 1.00 | -0.38 | -0.40 | 0.12 |
| *rsfS* | P73658 | Ribosomal silencing factor RsfS | -0.08 | 0.96 | 0.81 | -0.68 | 0.43 | -0.31 | -0.10 | 0.48 |
| *rpsP* | P74410 | Small ribosomal subunit protein bS16 | 0.45 | -0.14 | 0.06 | 0.61 | 0.19 | 0.00 | -0.96 | 0.20 |
| *aspS* | P73851 | Aspartate–tRNA(Asp/Asn) ligase | 0.08 | -0.67 | -0.55 | 0.20 | 0.28 | -0.58 | -0.96 | 0.16 |
| *rpsE* | P73304 | Small ribosomal subunit protein uS5 | 0.42 | -0.06 | -0.03 | 0.43 | 0.29 | 0.14 | -0.93 | 0.35 |
| *der* | P74120 | GTPase Der | 0.68 | -0.47 | -0.28 | 0.77 | 0.22 | -0.52 | -0.92 | -0.27 |
| *rplP* | P73313 | Large ribosomal subunit protein uL16 | 0.02 | -0.92 | -0.82 | -0.05 | 0.31 | 0.10 | -0.73 | 0.69 |
| *rpsS* | P73316 | Small ribosomal subunit protein uS19 | 0.41 | -0.32 | -0.59 | 0.91 | 0.14 | 0.10 | -0.65 | 0.40 |
| *rps1b* | P74142 | Small ribosomal subunit protein bS1B | 0.54 | 0.85 | 0.90 | 0.82 | 0.19 | 0.54 | -0.13 | 0.23 |
| *rpsC* | P73314 | Small ribosomal subunit protein uS3 | 0.12 | -0.45 | -0.40 | 0.26 | 0.52 | 0.33 | -0.51 | 0.89 |
| *rpsT* | P73336 | Small ribosomal subunit protein bS20 | 0.33 | -0.31 | -0.43 | 0.54 | 0.31 | 0.89 | 0.71 | 0.87 |
| *efp* | Q55119 | Elongation factor P | 0.06 | 0.89 | 0.52 | 0.00 | -0.11 | 0.55 | 0.24 | 0.26 |
| *rpsQ* | P73311 | Small ribosomal subunit protein uS17 | 0.18 | -0.06 | -0.10 | 0.55 | 0.00 | -0.26 | -0.88 | -0.08 |
| *rpsK* | P73298 | Small ribosomal subunit protein uS11 | 0.38 | -0.14 | 0.00 | 0.58 | 0.14 | -0.19 | -0.87 | -0.07 |
| *hisS* | Q55653 | Histidine–tRNA ligase | 0.01 | 0.01 | -0.56 | 0.37 | 0.03 | -0.86 | -0.53 | 0.10 |
| *cysS* | P74330 | Cysteine–tRNA ligase | 0.19 | 0.51 | 0.25 | 0.08 | -0.46 | -0.26 | -0.07 | -0.85 |
| *tuf* | P74227 | Elongation factor Tu | -0.18 | -0.64 | -0.82 | 0.19 | 0.06 | 0.22 | -0.76 | 0.19 |
| *pheS* | Q55187 | Phenylalanine–tRNA ligase alpha subunit | 0.29 | -0.07 | 0.54 | 0.18 | 0.56 | 0.07 | 0.78 | 0.81 |
| *rpmC* | P73312 | Large ribosomal subunit protein uL29 | 0.59 | 0.78 | 0.37 | 0.68 | -0.05 | 0.37 | 0.31 | -0.02 |
| *rplA* | P36236 | Large ribosomal subunit protein uL1 | 0.32 | 0.65 | 0.17 | 0.77 | -0.05 | -0.43 | -0.51 | -0.26 |
| *rpsN* | P48944 | Small ribosomal subunit protein uS14 | 0.09 | -0.53 | -0.40 | 0.76 | 0.19 | 0.11 | -0.72 | 0.56 |
| *infB* | P72689 | Translation initiation factor IF-2 | 0.14 | 0.30 | 0.02 | 0.49 | -0.14 | 0.05 | -0.75 | 0.49 |
| *rpmA* | P74267 | Large ribosomal subunit protein bL27 | 0.33 | -0.35 | -0.25 | 0.41 | 0.10 | 0.38 | -0.55 | 0.75 |
| *lepA* | P74751 | Elongation factor 4 | 0.35 | -0.13 | -0.26 | -0.13 | 0.18 | -0.75 | -0.23 | -0.08 |
| *rpsJ* | P74226 | Small ribosomal subunit protein uS10 | 0.06 | -0.47 | -0.45 | 0.42 | 0.20 | 0.03 | -0.73 | 0.32 |
| *infC* | P72874 | Translation initiation factor IF-3 | -0.05 | -0.59 | -0.52 | 0.63 | 0.33 | 0.29 | -0.56 | 0.72 |
| *rplJ* | P23350 | Large ribosomal subunit protein uL10 | 0.10 | 0.00 | -0.56 | 0.65 | 0.71 | -0.51 | 0.72 | -0.29 |
| *smpB* | P74355 | SsrA-binding protein | 0.60 | 0.59 | 0.14 | 0.25 | 0.12 | -0.20 | 0.71 | 0.00 |
| *rplS* | P36239 | Large ribosomal subunit protein bL19 | 0.32 | -0.21 | -0.18 | 0.71 | 0.23 | -0.17 | -0.60 | 0.09 |
| *fusB* | P74228 | Elongation factor G 2 | 0.28 | 0.53 | 0.69 | 0.45 | 0.09 | -0.30 | -0.11 | -0.13 |
| *rpmB* | P72851 | Large ribosomal subunit protein bL28 | 0.33 | -0.12 | -0.26 | 0.51 | 0.25 | 0.26 | -0.57 | 0.68 |
| *rpsH* | P73307 | Small ribosomal subunit protein uS8 | 0.20 | -0.23 | -0.11 | 0.32 | 0.37 | 0.56 | -0.53 | 0.67 |
| *rplD* | P73319 | Large ribosomal subunit protein uL4 | 0.15 | -0.53 | -0.16 | 0.38 | 0.32 | 0.41 | -0.65 | 0.25 |
| *ychF* | P73886 | Ribosome-binding ATPase YchF | 0.06 | -0.29 | -0.29 | 0.65 | 0.29 | 0.41 | -0.33 | 0.20 |
| *tsf* | P74070 | Elongation factor Ts | 0.03 | 0.04 | -0.22 | 0.46 | -0.07 | -0.11 | -0.57 | -0.05 |
| *serS* | P73201 | Serine–tRNA ligase | 0.07 | -0.01 | -0.27 | 0.21 | 0.06 | -0.29 | -0.26 | 0.57 |
| *bipA* | P72749 | Large ribosomal subunit assembly factor BipA | 0.18 | -0.46 | -0.43 | 0.57 | 0.32 | 0.04 | -0.51 | 0.33 |
| *thrS* | Q55806 | Threonine–tRNA ligase | 0.51 | 0.40 | 0.06 | 0.56 | 0.05 | -0.01 | -0.57 | -0.48 |
| *valS* | Q55522 | Valine–tRNA ligase | 0.12 | -0.16 | -0.38 | -0.07 | 0.18 | -0.51 | -0.56 | -0.15 |
| *rpsG* | P74229 | Small ribosomal subunit protein uS7 | 0.34 | -0.54 | 0.09 | 0.25 | 0.34 | -0.25 | 0.04 | 0.31 |
| *fusA* | P28371 | Elongation factor G 1 | 0.43 | -0.15 | 0.02 | 0.11 | 0.49 | 0.37 | 0.11 | -0.10 |
| *pheT* | P74296 | Phenylalanine–tRNA ligase beta subunit | -0.19 | 0.46 | 0.22 | 0.16 | 0.11 | 0.12 | -0.37 | 0.43 |
| *rpsO* | P72866 | Small ribosomal subunit protein uS15 | 0.28 | -0.26 | -0.32 | 0.29 | 0.40 | 0.05 | -0.42 | 0.11 |
| *rplN* | P73310 | Large ribosomal subunit protein uL14 | -0.06 | 0.20 | 0.07 | 0.09 | 0.14 | -0.20 | -0.41 | 0.03 |
| *asnS* | P52276 | Asparagine–tRNA ligase | 0.22 | -0.40 | -0.12 | -0.17 | 0.36 | -0.37 | -0.19 | 0.35 |
| **Purine & Pyrimidine Metabolism (n=13)** | | | | | | | | | | |
| *purU* | Q55135 | Formyltetrahydrofolate deformylase | 0.42 | -4.18 | -4.19 | -0.14 | 0.70 | 6.94 | 7.32 | 8.61 |
| *pyrH* | P74457 | Uridylate kinase | -1.06 | -3.58 | -3.02 | -0.38 | 0.68 | 0.28 | -0.66 | 1.28 |
| *pyrD* | P74782 | Dihydroorotate dehydrogenase (quinone) | -0.14 | -2.17 | -1.16 | 0.14 | -0.23 | -0.95 | -1.63 | 0.49 |
| *sll1459* | P73440 | Putative 5'-nucleotidase alr3139 | -0.30 | -1.19 | -1.39 | -0.54 | -0.13 | -1.59 | -2.11 | -0.82 |
| *pyrB* | P74163 | Aspartate carbamoyltransferase catalytic subunit | 0.79 | 1.51 | 1.54 | 1.77 | 0.44 | 0.58 | 0.50 | 0.50 |
| *slr0402* | P74432 | dITP/XTP pyrophosphatase | 0.05 | -0.97 | -0.25 | -0.02 | 0.47 | 0.34 | 1.22 | 1.51 |
| *prs* | Q55848 | Ribose-phosphate pyrophosphokinase | 0.05 | -1.36 | -0.87 | 0.37 | 0.20 | 0.66 | -0.56 | 0.54 |
| *pyrR* | Q55758 | Bifunctional protein PyrR | 1.18 | -0.08 | 1.33 | 0.76 | 0.94 | -0.78 | 0.19 | -0.56 |
| *purA* | P73290 | Adenylosuccinate synthetase | 0.38 | -0.22 | -0.47 | 1.13 | 0.38 | 0.14 | -1.06 | 0.21 |
| *purQ* | P73290 | Phosphoribosylformylglycinamidine synthase subunit PurQ | 0.38 | -0.22 | -0.47 | 1.13 | 0.38 | 0.14 | -1.06 | 0.21 |
| *nrdA* | P74240 | Ribonucleoside-diphosphate reductase subunit alpha | 0.60 | -0.32 | -0.19 | 0.68 | 0.53 | 0.28 | -0.48 | 1.07 |
| *upp* | P72753 | Uracil phosphoribosyltransferase | -0.32 | -0.15 | -1.05 | 0.44 | 0.16 | -0.20 | -0.97 | 0.00 |
| *pyrC'* | P72934 | Probable dihydroorotase-like protein | | | | | | | | |
